# Metabolic Adaptations To Acute Glucose Uptake Inhibition Converge Upon Mitochondrial Respiration For Leukemia Cell Survival

**DOI:** 10.1101/2024.11.20.624567

**Authors:** Monika Komza, Jesminara Khatun, Jesse D. Gelles, Andrew P. Trotta, Ioana Abraham-Enachescu, Juan Henao, Ahmed Elsaadi, Andriana G. Kotini, Cara Clementelli, JoAnn Arandela, Sebastian El Ghaity-Beckley, Agneesh Barua, Yiyang Chen, Bridget K. Marcellino, Eirini P. Papapetrou, Masha V. Poyurovsky, Jerry Edward Chipuk

## Abstract

One hallmark of cancer is the upregulation and dependency on glucose metabolism to fuel macromolecule biosynthesis and rapid proliferation. Despite significant pre-clinical effort to exploit this pathway, additional mechanistic insights are necessary to prioritize the diversity of metabolic adaptations upon acute loss of glucose metabolism. Here, we investigated a potent small molecule inhibitor to Class I glucose transporters, KL-11743, using glycolytic leukemia cell lines and patient-based model systems. Our results reveal that while several metabolic adaptations occur in response to acute glucose uptake inhibition, the most critical is increased mitochondrial oxidative phosphorylation. KL-11743 treatment efficiently blocks the majority of glucose uptake and glycolysis, yet markedly increases mitochondrial respiration via enhanced Complex I function. Compared to partial glucose uptake inhibition, dependency on mitochondrial respiration is less apparent suggesting robust blockage of glucose uptake is essential to create a metabolic vulnerability. When wild-type and oncogenic RAS patient-derived induced pluripotent stem cell acute myeloid leukemia (AML) models were examined, KL-11743 mediated induction of mitochondrial respiration and dependency for survival associated with oncogenic RAS. Furthermore, we examined the therapeutic potential of these observations by treating a cohort of primary AML patient samples with KL-11743 and witnessed similar dependency on mitochondrial respiration for sustained cellular survival. Together, these data highlight conserved adaptations to acute glucose uptake inhibition in diverse leukemic models and AML patient samples, and position mitochondrial respiration as a key determinant of treatment success.

## INTRODUCTION

Under normal conditions, cells obtain glucose from the bloodstream through facilitated diffusion by a family of glucose transporters (GLUT). There are 14 human GLUT proteins each with various tissue-specific expression patterns as well as differing substrate specificities and affinities [1, 2]. For example, GLUT1 and GLUT3 are the main transporters to shuttle glucose across the plasma membrane, while GLUT5 is selective for fructose [3, 4]. Once transported, glucose is phosphorylated by hexokinase, which keeps the cellular glucose concentration low and allows for continued uptake [5]. Glucose catabolism provides diverse carbon intermediates for macromolecule biosynthesis and enables sufficient production of the cofactors nicotinamide adenine dinucleotide (NADH) and flavin adenine dinucleotide (FADH_2_) via the tricarboxylic acid (TCA) cycle. These cofactors couple glycolysis to mitochondrial respiration and adenosine triphosphate (ATP) production [6]

Rapidly proliferating cells exhibit a high demand for glucose; and in the case of cancer cells, increased glycolysis occurs despite aerobic conditions, a phenomenon known as the Warburg effect [7]. Increased glycolysis is a key feature of almost all cancers, including haematological malignancies (*e.g.,* acute myeloid leukemia (AML)), and is a prime example of metabolic adaptations that enable cancer cells to meet the high bioenergetic demands of hyper-proliferation [8, 9]. Oncogenic signalling pathways (*e.g.,* RAS^G12D^) promote glucose uptake by increasing the transcription and plasma membrane localization of GLUT1 and GLUT3 along with glycolytic enzymes [10–13]. Moreover, a glycolytic metabolic signature identified in AML patients indicates increased glycolytic flux correlates with poor survival outcomes, and increased glycolysis is also associated with chemotherapy resistance [14].

Given the relationships between glucose metabolism and cancer, this pathway is an indisputable pharmacological target. Inhibitors of glycolytic enzymes were developed as potential therapeutics, but they act downstream of the growth factor-independent glucose uptake observed in many cancers [15–18]. As glycolytic intermediates often shuttle to various metabolic pathways supporting tumor growth, a more direct alternative approach has emerged in the past decade focused on glucose uptake inhibition [19–21]. Several GLUT1 inhibitors, including WZB117, STF-31, NV-5440, and BAY-876 are described, but while GLUT1 is the primary transporter affected by cancer signaling, additional glucose transporters promote tumor cell survival and often gain function upon GLUT1 inhibition – all of which provides a rationale for targeting multiple GLUTs [22–25]. Most recently, Kadmon Corporation developed a pan Class I GLUT1-4 inhibitor (KL-11743) through a cell-based phenotypic assay assessing inhibition of non-mitochondrial ATP production followed by structure-activity relationship optimization [26]. While KL-11743 effectively blocks glucose uptake in the nanomolar range, subsequent cellular phenotypes and metabolic adaptions remain largely unknown thus limiting this strategy’s translational potential.

In this study, we systematically characterized metabolic responses to KL-11743 using a panel of leukemia cells of similar origin but with varying metabolic and genetic backgrounds. We hypothesized that KL-11743 would effectively inhibit glycolysis in all leukemia models, yet this would be insufficient to promote a chemotherapeutic response given the likelihood of metabolic adaptations. Indeed, we demonstrate that KL-11743 effectively inhibits glucose uptake and reveals a bioenergetic vulnerability, which is exploited by NADH dehydrogenase inhibition; importantly, these responses are conserved from cell lines, patient stem-cell models of leukemia, and AML patient samples. Our work underscores the complex metabolic landscape and plasticity of cancer cells and proposes a general model of mapping mitochondrial responses to acute GLUT1-4 inhibition.

## METHODS

### Reagents

Standard reagents were obtained from ThemoFisher Scientific (MA, USA) and Sigma-Aldrich (MO, USA). KL-11743 and KL-12023 were kindly provided by Kadmon Pharmaceuticals (NY, USA). Oligomycin, carbonyl cyanide-p-trifluoromethoxyphenylhydrazone (FCCP), rotenone, antimycin A, aminooxyacetic acid, etomoxir, cycloheximide, phenylsuccinic acid, 2-(aminooxy)acetic acid, 3-nitropropionic acid, and thenoyltrifluoroacetone were from Sigma-Aldrich. ABT-737, IACS-010579, and GSK-1120212 were from Selleck Chemicals (TX, USA). MitoTracker^TM^ Green FM, Tetramethylrhodamine ethyl ester, and DAPI were from ThermoFisher Scientific. All primary cell culture additives were from Stem Cell Technologies (MA, USA).

### Cell Models

NB4, THP-1, and MOLT-4, lines were obtained from American Type Cell Culture. Cells were cultured in Roswell Park Memorial Institute Medium (RPMI-1640) containing phenol red and 300 mg/L L-glutamine (ThermoFisher Scientific, MA, USA) and supplemented with 10% heat inactivated fetal bovine serum, 100 U/mL penicillin, and 100 ug/mL streptomycin (ThermoFisher Scientific, MA, USA). Glutamine deficient, serine deficient and glutamine/serine double deficient RPMI media were purchased from Memorial Sloan-Kettering Cancer Center Media Preparation Facility. Cells were incubated in a humidified incubator at 37°C with 5% CO_2_ and confirmed to be mycoplasma-free by the HEK-Blue Detection Kit (Invitrogen, MA, USA). Induced pluripotent stem cell (iPSC) lines encompassing normal cells (N-8.2), KRAS^WT^ AML (AML-4.24), and KRAS^G12D^ (AML-4.10) AML were generated, cultured and differentiated into HSPCs/LSCs as previously described[27, 28]. Patient AML samples were cultured in RPMI-1640 supplemented with 10% fetal calf serum, 1% penicillin, 1% streptomycin, 2 mM glutamine, 5 μM β-mercaptoethanol, 20 ng/ml hIL-3, 50 ng/mL hIL-6, 20 ng/ml hGM-CSF, 20 ng/ml hG-CSF, 20 ng/ml hTPO, and 25 ng/mL hSCF. De-identified patient samples were provided by the Tisch Cancer Institute’s Hematological Malignancies Tissue Bank through an Institutional Review Board at the Mount Sinai School of Medicine approved protocol (STUDY-11-02054-MOD008; PI: Bridget Marcellino).

### Glucose Uptake Assays

Cells were seeded in 96-well plates in quadruplicate and treated with DMSO or KL-11743 (500 nM) for 24 h. The day of the assay, 200 μL was removed and centrifuged at 400 × *g* for 5 min. Resulting cell pellets were resuspended in fresh media, transferred back to their respective wells, and normalized by YOYO3 staining, described below. Supernatant was used to measure glucose content using the Glucose (GO) assay kit (Sigma-Aldrich) scaled down to be assessed on a 96-well tissue culture plate and performed according to the manufacturer’s instructions. Absorbance readings were measured at 540 nm using a plate reader (Synergy H1 Hybrid multi-mode microplate reader, Biotek/Agilent, CA, USA). Concentrations were determined by a standard curve, then normalized to cell count and calculated as a percentage relative to base glucose levels in culture media containing 10% FBS.

### Nutrient Consumption Analysis

Cells were treated with KL-11743 (500 nM) or DMSO. At 24 h, glucose, lactate, glutamine, and glutamate concentrations in the medium were measured with an YSI7000 electrochemical analyzer (YSI) in collaboration with the Donald B. and Catherine C. Marron Cancer Metabolism Center at Memorial Sloan Kettering Cancer Center (NY, USA).

### Amino Acid Consumption Analysis

Cells were treated with KL-11743 (500 nM) or DMSO for 24 hours and amino acid concentrations were determined by YSI7000 electrochemical analyzer (YSI) in collaboration with Mount Sinai Metabolomics Core. For metabolite measurements from spent culture medium, 50 μL of cell-conditioned medium was extracted by the addition of 200 μL of 80:20 ice-cold methanol:water and stored at −80°C overnight. 50 μL of blank medium incubated for the same amount of experimental time was processed in parallel and used as a reference to determine metabolite secretion or consumption. The methanol-extracted metabolites were cleared by centrifugation at 20,000g for 20 minutes at +4°C, and supernatant was dried in a vacuum evaporator (Genevac EZ-2 Elite) for 2 hours. Dried metabolites were dissolved in 20 mg/mL of methoxyamine hydrochloride (Sigma-Aldrich, 226904) in pyridine (Thermo Fisher, TS-27530) for 90 min at 30°C and derivatized with MSTFA with 1% TMCS (Thermo Fisher, TS-48915) for 30 min at 37°C. Samples were analyzed using an Agilent 7890A GC connected to an Agilent 5975C Mass Selective Detector with electron impact ionization.

### Normalization for Cell Count

Cells were treated with 0.1% Triton and 0.125 μM YOYO™-3 Iodide (Invitrogen, MA, USA) and incubated at room temperature overnight, protected from light. 599/640 nm (excitation/emission) wavelength was measured with Synergy H1 Hybrid multimode microplate reader (BioTek/Agilent). Fluorescence unit values were divided by the 20% trimmed mean calculated from all wells to obtain normalization factors.

### Cell Counts For Proliferation & G1-Phase Cell Cycle Arrest Analysis

Cells were plated at an initial concentration of 2×10^5^ cells/mL and treated with 500 nM KL-11743 or an equal volume of DMSO. Every 12 h, a portion of the cell suspension was removed, pelleted, resuspended in a smaller volume, and counted using a hemocytometer. Trypan Blue Solution (0.4%) (ThermoFisher Scientific) was used at a 1:1 ratio to stain and exclude dead cells. For G1-phase cell cycle arrest quantification, cells were treated as indicated, trypsinized, washed with PBS, and resuspended in Nicoletti-Buffer (0.1% TX-100, 0.1% sodium citrate, 50 μg/ml propidium iodide). Intact nuclei were analyzed by flow cytometry to quantify G1 DNA content. Data analysis was conducted using FCS Express 7 computer software. Unless indicated otherwise, flow cytometry data was collected by the Flow Cytometry CoRE at the Icahn School of Medicine at Mount Sinai.

### EdU Flow Cytometry Assays

Percentages of cells proliferating were obtained using the Click-iT™ EdU Alexa Fluor™ 647 Flow Cytometry Assay Kit (Invitrogen) and prepared according to manufacturer’s instructions. Briefly, cells were treated as indicated for 24 h, followed by a 2 h incubation with 10 μM EdU at 37°C. Cells were harvested, washed, permeabilized, and fixed, and analyzed by flow cytometry at 633/635 nm excitation. Data analysis was conducted using FCS Express 7 computer software. Unless indicated otherwise, flow cytometry data was collected by the Flow Cytometry CoRE at the Icahn School of Medicine at Mount Sinai.

### Apoptosis Assays

Cell lines were treated in 12-well plates as indicated, with at least 3×10^5^ cells per sample. The day of the assay, cell suspensions were transferred to 12×75 mm polystyrene flow cytometry tubes, pelleted at 1000 × *g* for 10 min at 4°C and resuspended in 300 μL Annexin-V Binding Buffer (10 mM HEPES pH 7.4, 150 mM NaCl, 5 mM KCl, 1 mM MgCl_2_, 1.8 mM CaCl_2_) containing ∼3.5 μg/μl AlexaFluor 488-conjugated Annexin V. Samples were analyzed for percent positive events by flow cytometry. Data analysis was conducted using FCS Express 7 software. Unless indicated otherwise, flow cytometry data was collected by the Flow Cytometry CoRE at the Icahn School of Medicine at Mount Sinai. Patient AML samples were treated as indicated and cell death was measured using an MTT assay (Sigma-Aldrich) according to the manufacturer’s instructions. This minimized harvesting/labelling stress and processing times to provide more accurate datasets compared to Annexin V analysis of primary cells. Sample were analyzed by measuring the A590 and subtracting the A690 as a reference; % viability was calculated by normalizing absorbance values to DMSO (the negative control) and cycloheximide (the positive control) to 0 and 100%, respectively.

### Viability Assays

Cells were grown in complete RPMI, gln deficient RPMI, ser deficient RPMI, and gln/ser double deficient RPMI media with DMSO or KL-11743 treatment as indicated in a 96-well plate at 3×10^4^ cells per well. CellTiter-Glo 2.0 (Promega) was used to measure the viability after 24 hours following manufacturer’s protocol. Readings were taken in Agilent BioTek Synergy H1 plate reader.

### Single-Cell and Population-Level Analyses Using Real-Time Kinetic Labeling (SPARKL)

Cells were seeded onto poly-D-lysine (ThermoFisher Scientific)-coated 96-well plates at 3×10^4^ cells per well. Cells were treated as indicated in addition to 1 μM cell viability dye YOYO™-3 Iodide (Invitrogen) before immediately subjecting the plate to real-time cell-death analysis. Cells were incubated in a humidified and gas-controlled environment and imaged using a tandem BioSpa and Cytation 7 Cell Imaging Multimode Reader (BioTek/Agilent) as described previously[29]. A 1043 × 1043 μm image was taken per well, analyzed for fluorescently positive objects, and data is reported as the number of positive objects detected per image. Images were collected using a 10× objective, a laser auto focus module (cat. no. 1225010), and the “Texas Red” filter cube – excitation: 586/15, emission: 647/57 (cat. no. 1225102, BioTek/Agilent).

### NAD^+^/NADH Analyses

NAD^+^ and NADH concentrations were determined using the EnzyChrom™ NAD^+^/NADH Assay Kit (BioAssay Systems, CA, USA). 1×10^6^ cells were treated at a concentration of 2×10^5^ cells/mL. Cells were pelleted, resuspended in NAD^+^ or NADH extraction buffer, and analyzed following manufacturer’s instructions. Optical density was read for time “zero” at 520–600 nm and again after a 15 min incubation at room temperature. Measurements from time “zero” were subtracted from the final reads, and NAD^+^/NADH concentrations were determined by plotting measurements against a standard curve of known NAD concentrations.

### Targeted Metabolomics

NB4 (Fig. S2): NB4 was treated with DMSO or KL-011743 (500 nM, 24 h), pelleted, and frozen at −80°C. Samples was thawed on ice, 100 μL of ultrapure water was added to resuspend the cell pellet. Divide 50 μL cell suspension and add 200 μL of methanol (precooled at −20°C) and vortexed for 2 min under the condition of 2500 r/min. The sample was frozen in liquid nitrogen for 5 min, removed on ice for 5 min, after that, the sample was vortexed for 2 min. The previous step was repeated for 3 times. The sample was centrifuged at 12000 r/min for 10 min at 4°C. Take 200 μL of supernatant into a new centrifuge tube and place the supernatant in −20°C refrigerator for 30 min. Then the supernatant was centrifuged at 12000 r/min for 10 min at 4°C. The sample extracts were analyzed using an LC-ESI-MS/MS system (Waters ACQUITY H-Class, https://www.waters.com/nextgen/us/en.html; MS, QTRAP^®^ 6500+ System, https://sciex.com) by Metware Biotechnology. Metabolites were quantified by multiple reaction monitoring (MRM) using triple quadrupole mass spectrometry. In MRM mode, the first quadrupole screened the precursor ions for the target substance and excluded ions of other molecular weights. After ionization induced by the impact chamber, the precursor ions were fragmented, and a characteristic fragment ion was selected through the third quadrupole to exclude the interference of non-target ions. After obtaining the metabolite spectrum data from different samples, the peak area was calculated on the mass spectrum peaks of all substances and analyzed by standard curves. Profiling Of Widely Targeted Small Metabolites By QTRAP 6500+ LC-MS/MS (Figs. 3A–D, S4A): NB4 and MOLT4 were treated with DMSO or KL-011743 (500 nM, 24 h), pelleted, washed with 1 mL of 150 mM ammonium acetate, pelleted again, and flash-frozen in liquid nitrogen. Samples were screened for alterations in metabolite levels of the Widely Targeted Small Polar Metabolite (WTSM) panel by the Stable Isotope and Metabolomics Core Facility at Albert Einstein College of Medicine, as previously described [30].

### KEGG Pathway Enrichment Analysis

Rich Factor for each pathway, the ratio of the number of differential metabolites in the corresponding pathway to the total number of metabolites annotated in the same pathway, was calculated. The greater the Rich Factor, the greater the degree of enrichment. P-value is the calculated using hypergeometric test as shown below:

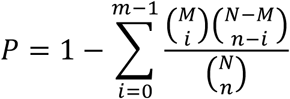

N represents the total number metabolites with KEGG annotation, n represents the number of differential metabolites in N, M represents the number of metabolites in a KEGG pathway in N, and m represents the number of differential metabolites in a KEGG pathway in M.

### Real-Time Reverse Transcription Polymerase Chain Reaction

Cells were treated as indicated for 24 h, then harvested and pelleted by centrifugation for 5 min at 400 × *g*. Total RNA was extracted from cell pellets with RNeasy Mini Kit (Qiagen, MD, USA), following manufacturer’s instructions. RNA was quantified using a NanoDrop One Spectrophotometer, and 2 μg was used to synthesize cDNA using the RNA to cDNA EcoDry^TM^ PreMix (Double Primed) (Takara Bio, CA, USA). Reaction tubes were placed in a thermocycler and subjected to 42°C for 1 h and 70°C for 10 min, followed by an indefinite 4°C hold until moved to storage at −20°C. Forward and reverse primers for genes of interest (Table 1) were combined with *Power* SYBR™ Green PCR Master Mix (Applied Biosystems, CA, USA), and gene expression was analyzed using a ViiA 7 Real-Time PCR system. The expression of relevant genes was normalized to *18S*.

**Table 1.**
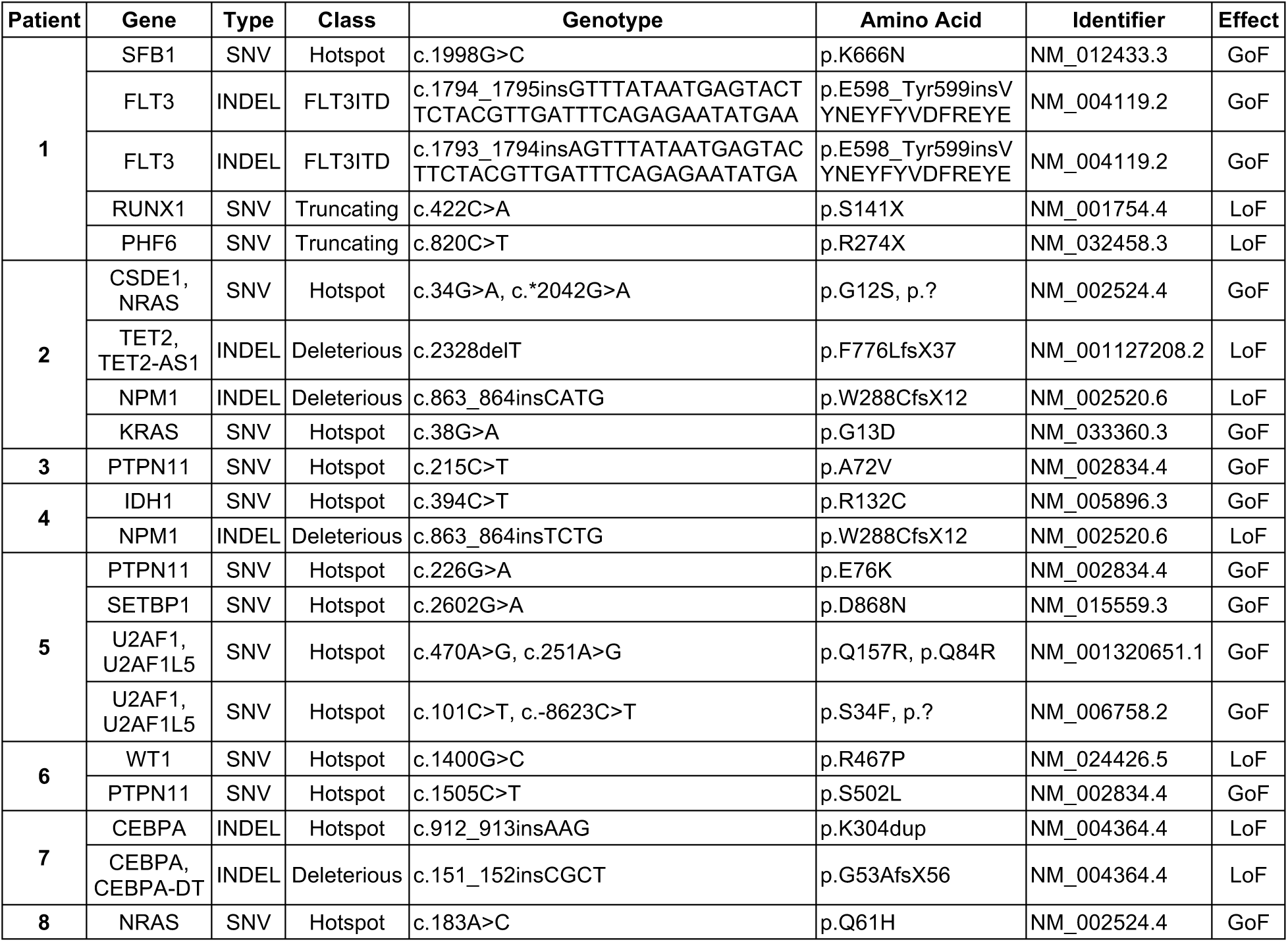
Primary AML patient samples mutational status.

**Table 2.**
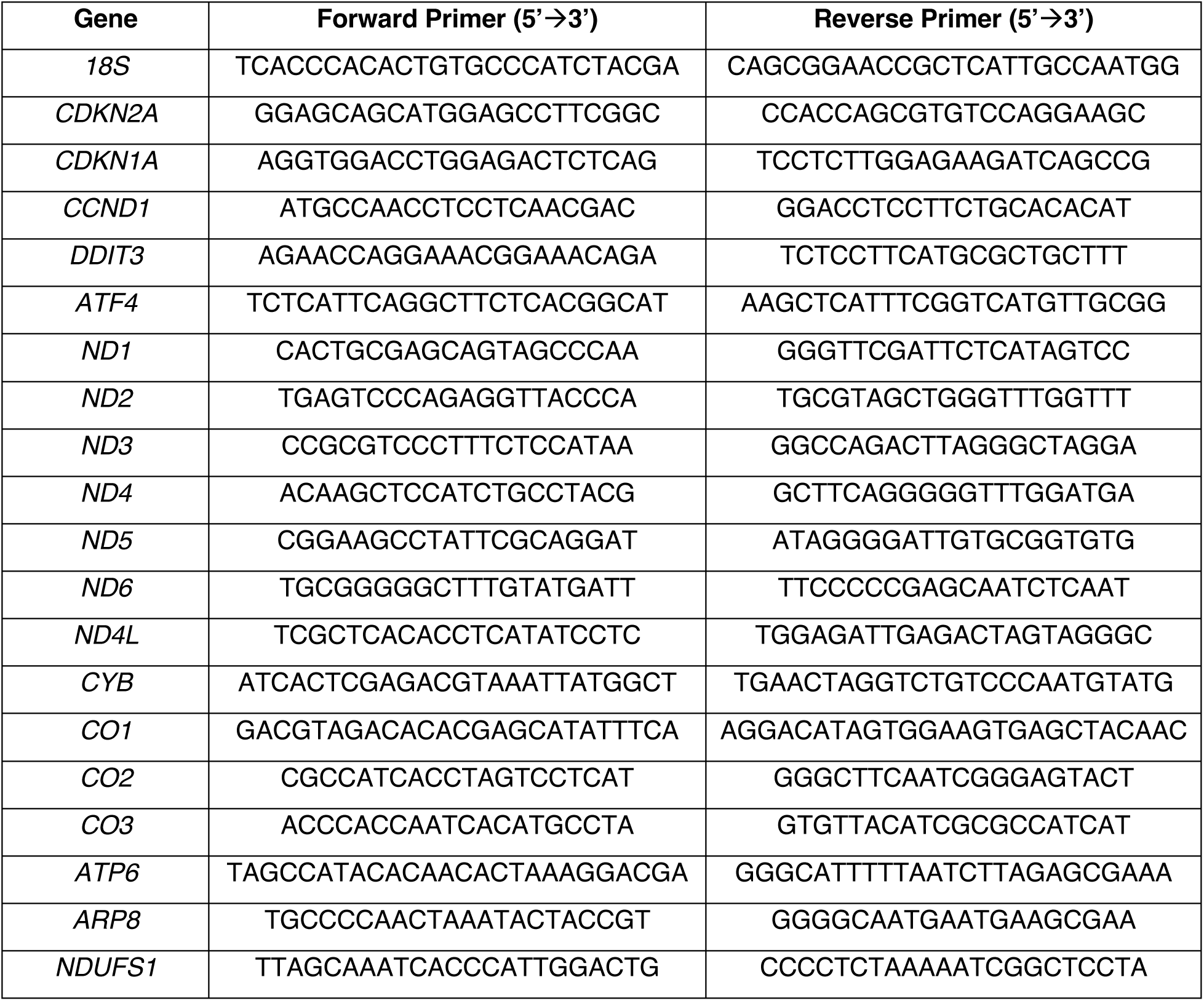
qPCR primer sequences for probed genes.

### Agilent Bioanalyzer Glycolysis Stress Test Analysis

Cells were treated as indicated at a concentration of 2×10^5^ cells/mL. One day prior to running the assay, Agilent XFe96 Sensor Cartridges were hydrated with XF Calibrant, pH 7.4 (Agilent). The day of the assay, cells were pelleted and resuspended in XF RPMI (Agilent) supplemented with 2 mM L-glutamine and the indicated treatment condition. Cells were counted and seeded (1×10^5^ cells/well) onto poly-D-lysine (ThermoFisher Scientific)-coated plates (Agilent) and centrifuged at 200 × *g* for 3 min. Plates were incubated in a non-CO_2_ incubator at 37°C for 40–60 min. OCR and ECAR were measured using the Agilent XFe96 Extracellular Flux Analyzer and the XF Glycolysis Stress Test Kit (Agilent) according to the manufacturer’s instructions. ECAR measurements were determined before and after administration of glucose (10 mM), oligomycin (1 μM), and 2-deoxy-D-glucose (50 mM). At the end of the assay, normalization values for cell count were obtained as described above (*“Normalization for Cell Count”*), and OCR and ECAR measurements were normalized against these factors.

### Agilent Bioanalyzer XF Cell Mito Stress Test

Cells were treated as indicated at a concentration of 2×10^5^ cells/mL. One day prior to running the assay, Agilent XFe96 Sensor Cartridges were hydrated with XF Calibrant, pH 7.4 (Agilent). The day of the assay, cells were pelleted and resuspended in XF RPMI (Agilent) supplemented with 1 mM pyruvate (Agilent), 10 mM glucose (Agilent), 2 mM L-glutamine, as well as the treatment condition. Cells were counted and seeded (1×10^5^ cells/well) onto poly-D-lysine-coated plates (Agilent) and centrifuged at 200 × *g* for 3 min. Plates were incubated in a non-CO_2_ incubator at 37°C for 40–60 min. OCR and ECAR were measured using the Agilent XFe96 Extracellular Flux Analyzer and the Agilent XF Cell Mito Stress Test (Agilent) according to the manufacturer’s instructions. OCR was measured 3 times before and after administration of each of the following: oligomycin (1 μM), FCCP (1 μM), and a combination of rotenone and antimycin A (0.5 μM). At the end of the assay, normalization values for cell count were obtained as described above, and OCR and ECAR measurements were normalized against these values.

### Agilent Bioanalyzer Complex I/II Activities

Cells were treated as indicated at a concentration of 2×10^5^ cells/mL. One day prior to running the assay, Agilent XFe96 Sensor Cartridges were hydrated with XF Calibrant, pH 7.4. Cells were pelleted and resuspended in mitochondrial assay buffer (MAS: 220 mM mannitol, 70 mM sucrose, 10 mM KH_2_PO_4_, 5 mM MgCl_2_, 2 mM HEPES, 1 mM EGTA, 0.2% fatty acid-free BSA). Cells were counted and seeded (1×10^5^ cells/well) onto poly-D-lysine-coated plates (Agilent), centrifuged at 200 *× g* for 3 min, and incubated at 37°C for 30 min. 100 μL MAS was removed from each well and replaced with 100 μL MAS containing plasma membrane permeabilizer for a final concentration of 1 nM in the well. Sensor Cartridge was loaded with 10× stocks of substrates and/or inhibitors diluted in MAS without BSA. For CI analysis, CI was stimulated with pyruvate (10 mM), malate (0.5 mM), and ADP (4 mM); CI was inhibited with rotenone (1 μM); CII was stimulated by succinate (10 mM); and maximal respiration was stimulated with FCCP (1 μM). For CII analysis, CI was inhibited with rotenone (1 μM) and CII stimulated with succinate (10 mM) and ADP (4 mM); ATP-linked respiration was inhibited with oligomycin (1 μM); maximal respiration was stimulated by FCCP (1 μM); and respiration was decreased to non-mitochondrial levels with antimycin A (0.5 μM). At the end of the assay, normalization values for cell count were obtained as described above (*“Normalization for Cell Count”*), and OCR measurements were normalized against these values.

### Heavy Membrane Isolations

Cells (∼6×10^7^ per sample) were treated as indicated for 24 h. Cells were then pelleted at 400 × *g* for 5 min at 4°C and washed twice with 3 mL trehalose isolation buffer (TIB: 300 mM trehalose, 10 mM HEPES, 10 mM KCl, 1 mM EGTA, 0.1% BSA fraction V). Cells were resuspended in TIB supplemented with protease inhibitor and kept on ice. Cell suspensions were passed through a 27-gauge needle 13 times. The homogenate was centrifuged at 1,000 × *g* at 4°C for 10 min, the supernatant collected, and centrifuged again at the same conditions to ensure that no unlysed cells or nuclei were present. The resulting supernatant was centrifuged at 10,000 *× g* at 4°C for 10 min, and the resulting pellet collected as the heavy membrane isolate.

### In-Gel Mitochondrial Complex Extractions and Analyses

Heavy membranes were resuspended in extraction butter (1 M 6-amino-hexanoic acid, 50 mM Bis-Tris HCl pH 7.0) and quantified using a Pierce™ BCA Protein Assay Kit (ThermoFisher Scientific) according to manufacturer’s instructions. 90 μg was solubilized with digitonin (final concentration 6%) on ice for 10 min, followed by centrifugation at 13,000 × *g* at 4°C for 20 min. Supernatant was collected and glycerol added for a final concentration of 10%. 45 μg of sample/lane was resolved using Criterion TGX^TM^ 4%-15% gels (Bio-Rad Laboratories, CA, USA) using native conditions at 50 V for about 4 h. For in-gel CI assays, gels were soaked in 30 mL of 5 mM Tris/HCl pH 7.4 containing NTB (30 mg) and 300 μL of 10 mg/mL NADH; for in-gel CII assays, gels were soaked in 30 mL of 5 mM Tris/HCl pH 7.4 containing NTB (30 mg), 600 μL of 1 M sodium succinate, and 24 μL of 250 mM phenazine methosulfate. All in-gel assays were performed for 1 h at room temperature. 15 μg of samples were resolved on a separate Criterion TGX^TM^ 4%-15% gel under native conditions at 120 V and stained with Coomassie Blue as a loading control. Bands were quantified using (Fiji Is Just) ImageJ.

### Mitochondrial Mass and Mitochondrial Membrane Potential Analyses

Cells were plated at an initial concentration of 2×10^5^ cells/mL and treated with 500 nM KL-11743 or an equal volume of DMSO for 24 h. Untreated parental cells were incubated without fluorescent staining or FCCP treatment for an unstained control. FCCP was used as a positive control for both membrane depolarization and accumulation of mitochondrial ROS for each cell line by treating with 10 μM FCCP before placing in the dark at 37°C with 5% CO_2_ for 30 min. Cells were stained for 30 min using 200 nM MitoTracker^TM^ Green FM or 100 nM TMRE, supplemented with DMSO, KL-11743, or FCCP, and placed in the dark at 37°C with 5% CO_2._ Cells were collected in round-bottom polypropylene test tubes and centrifuged for 5 min at 450 × *g* at 4°C. Supernatants were discarded and pellets were resuspended with 1 μg/mL DAPI in 300 μL of 1× PBS, supplemented with DMSO, KL-11743, or FCCP. Unstained parental cells were resuspended in 300 μL of 1× PBS without DAPI. All prepared stained cells and controls were analyzed with a BD FACSCanto II Clinical Flow Cytometry System with the parameters defined by the unstained and stained parental cells. All conventional flow cytometry data analysis was conducted through the FCS Express 7 computer software (version: 7.14.0020). An initial side scatter-area (SSC-A) vs forward scatter-area (FSC-A) density plot was used to gate on the population based on size and granularity while removing debris and dead cells found on the bottom left corner of the density plot due to a lower level of FSC-A. To exclude doublet cells from the population of interest, a second gate was established on the population from former gate by plotting FSC-height (FSC-H) vs FSC-A and identifying disproportions in area. To effectively discriminate dead cells in the population, DAPI viability staining, excitable by the 405 nm laser (Violet-1), was used, and FSC-A vs Violet-1 density plots were generated to exclude DAPI positive cells. The population of interest was portrayed using a final density plot with the y-axis set to FSC-A and the x-axis set to the fluorophore’s channel. Data export and analysis was performed on the population from this final density plot. Overall cell size was considered upon analysis of mitochondrial mass and mitochondrial membrane potential by flow cytometry as raw FSC-A values for each cell was divided by the mean FSC-A of the population of interest to convert FSC-A values to a normalization factor centered on 1.00. Mitochondrial mass and mitochondrial membrane potential were then determined by normalizing each cell’s mean fluorescent intensity (MFI) for MitoTracker^TM^ Green FM or TMRE against the normalized FSC-A values.

### Genomics Analyses

DNA sequencing: AML patient’s peripheral blood mononuclear cells were employed for extracting genomic DNA using a DNeasy Blood & Tissue Kit (Qiagen) following the manufacturer’s instructions. DNA quality and quantity were assessed by Agilent 2100 Bioanalyzer system and Qubit (ThermoFisher Scientific). DNA sequencing and mutational analyses were performed using the Ion Torrent platform and Ion Reporter v5.14; annotations were defined using the Oncomine Myeloid Assay Annotations v1.2 r.0, and using the following references: hg19, Oncomine Myeloid DNA Hotspots v1.3, Oncomine Myeloid DNA Mask – 318 – v1.2, Oncomine Myeloid DNA Regions – 318 – v1.0 (ThermoFisher Scientific). Cell line RNA-Seq: NB4, and THP-1, RNA-seq scaled mRNA expression was performed using The Broad Institute’s Cancer Cell Line Encyclopedia (www.broadinstitute.org/ccle). iPSC RNA-Seq: Analysis was performed on the publicly available data GEO: GSE92494. The RNA read counts and metadata were obtained using the function “getGEO” from the R-package “GEOquery”. This data was later rearranged to construct a “DESeqDataSet” object containing only the samples of interest and genes with more than 10 reads. Principal component analysis was conducted using the function “plotPCA” on the regularized log transformation (“rlog” function of the R-package “DESeq2”) of normalized read counts. The differential expression analysis was performed with the R-package “DESeq2” comparing normal versus AML samples, yielding as significant those with an adjusted *p*-value (Benjamini & Hochberg) <0.05 and log2FC > 2 or log2FC < −2. To generate the heatmap of 1460 differentially expressed genes, the read counts were scaled, log normalized and plotted using the R-package “ComplexHeatmap” with default row clustering. To visualize ontology pathways of interest, the significance cutoff was set at an adjusted *p*-value (Benjamini & Hochberg) <1E-5 and log2FC > 1 or log2FC < −1. The differentially expressed data frame was filtered by each gene list downloaded from the Molecular Signatures Database (MSigDB) and graphed as volcano plots using Prism - GraphPad and Inkscape. The Reactome Pathway Database (Mitochondrial Protein Import and Translation), The Hallmark gene sets (Oxidative Phosphorylation and Glycolysis), and WikiPaths (Electron Transport Chain assembly) were utilized.

## RESULTS

### KL-11743 inhibits glucose uptake and stimulates mitochondrial respiration

To evaluate baseline glucose uptake in our leukemia cell models, we measured the percentage (%) of glucose consumed by NB4, THP-1, and MOLT-4 cells over a 24 h period. MOLT-4 cells exhibited the highest glucose consumption (∼97%), followed by NB4 (∼86%) and THP-1 (∼63%) cells (Fig. 1A). Treatment with KL-11743 for 24 h effectively inhibited glucose uptake levels in a dose-dependent manner (Figs. 1B–D, S1A–C). We investigated the possibility of an irreparable metabolic deficiency by assessing proliferation rates. In NB4 cells, a noticeable difference in cell counts was observed as early as 12 h post-treatment (Fig. 1E). THP-1 and MOLT-4 cells, which proliferate slower than NB4 cells, did not exhibit a discernible decrease in cell counts until 24–36 h post-treatment (Figs. 1F–G). To quantify differences in the percent of cells actively proliferating at the 24 h timepoint, we stained with EdU. We found that NB4 cells showed the largest decrease in proliferating cells upon KL-11743 treatment, while THP-1 and MOLT-4 cells exhibited a more modest decrease (Fig. 1H). To further analyze proliferation at the 24 h timepoint, we examined mRNA expression of cyclin-dependent kinase inhibitors *CDKN2A* (p16) and *CDKN1A* (p21), as well as mRNA expression of *CCND1* (cyclin D). We observed a modest increase in *CDKN1A* expression in NB4 and MOLT-4 cells, suggesting some influence on cell cycle; however, cyclin D expression was largely unaltered, indicating that KL-11743-treated cells continue to cycle and proliferate, albeit at slower rates (Fig. 1I).

**Figure 1.**
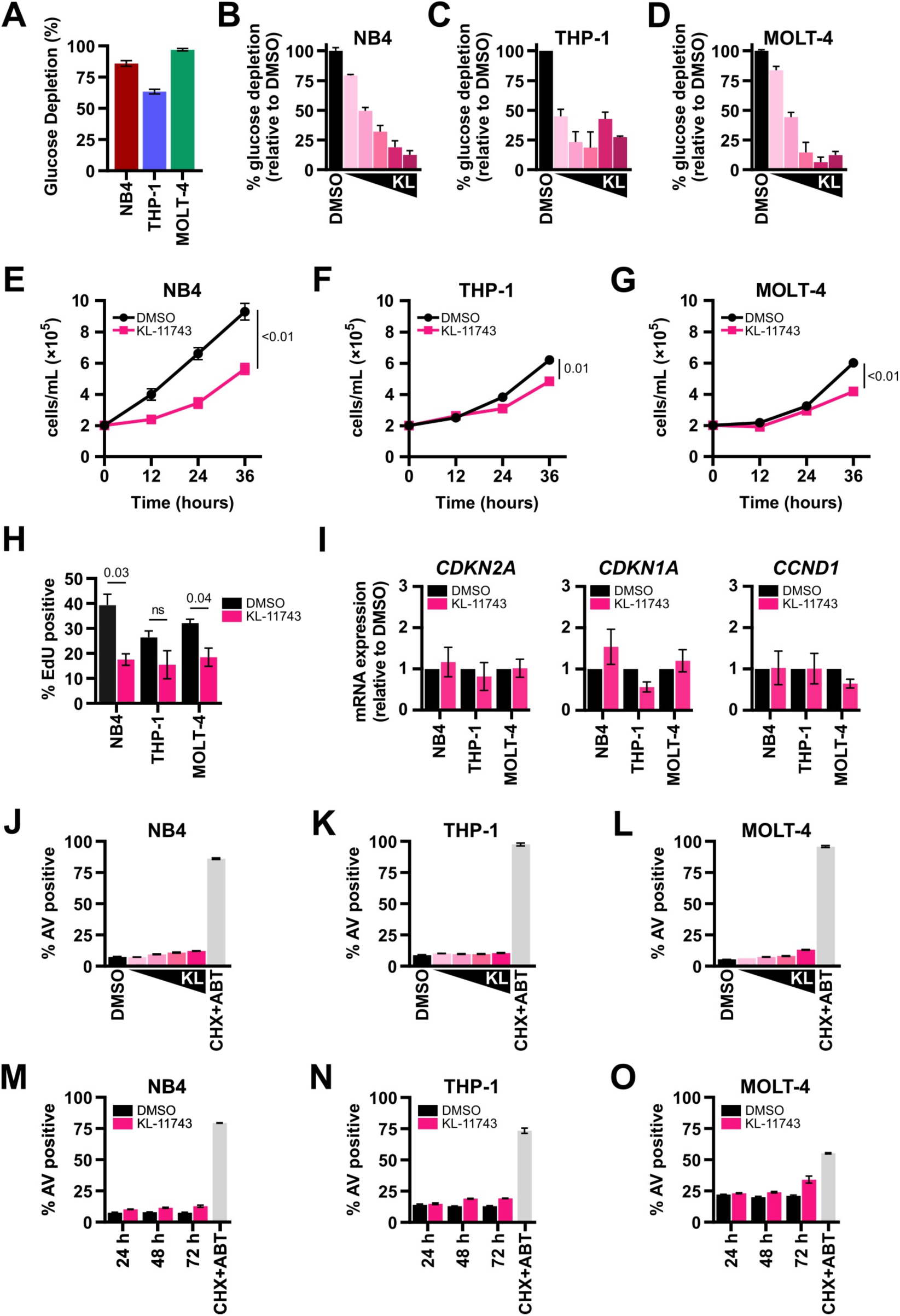
KL-11743-mediated inhibition of glucose transporters reduces glucose uptake and proliferation rates but does not induce cell death. **(A)** Glucose concentrations were assessed in conditioned media and unconditioned RPMI for the indicated cell lines at 24 h. The difference was calculated to obtain % glucose depletion. **(B–D)** NB4, THP-1, and MOLT-4 were treated with DMSO or KL-11743 (50, 150, 500, 1500, 2500 nM) for 24 h. Glucose concentrations were assessed in conditioned media and unconditioned RPMI, and the difference was calculated to obtain glucose depletion. Data are presented as % depletion relative to DMSO. **(E–G)** NB4, THP-1, and MOLT-4 were treated with DMSO or KL-11743 (500 nM) and cells were manually counted every 12 h, excluding dead cells as identified by staining with Trypan Blue. **(H)** NB4, THP-1, and MOLT-4 were treated with DMSO or KL-11743 (500 nM) for 24 h. Cells were harvested, treated with EdU (10 μM) for 2 h, fixed, and analyzed by flow cytometry. **(I)** NB4, THP-1, and MOLT-4 were treated with DMSO or KL-11743 (500 nM) for 24 h, and total RNA was harvested. The fold change of transcripts for p16 (*CDKN2A*), p21 (*CDKN1A*), and cyclin D (*CCND1*) were measured by real-time qPCR. Expression was normalized against *18S*. **(J–L)** NB4, THP-1, and MOLT-4 were treated with DMSO or KL-11743 (10, 100, 250, 500 nM) for 24 h. Apoptosis was measured by Annexin V (AV) labeling and flow cytometry. Cycloheximide (CHX, 50 μg/mL) and ABT-737 (ABT, 1 μM) is a positive control for apoptosis. **(M–O)** NB4, THP-1, and MOLT-4 were treated with DMSO or KL-11743 (500 nM) for 24, 48, or 72 h and apoptosis was measured. All data are presented as mean values of at least 3 replicates ± SEM. Where indicated, *p* values were calculated using an unpaired T test.

In the literature, it is well-established that glucose limitation activates the endoplasmic reticulum (ER) unfolded protein response (ER^UPR^), and this pathway influences both cell cycle and cell death[31, 32]. To determine if KL-11743 activates the ER^UPR^, we treated NB4, THP-1, and MOLT-4 with KL-11743 for 8 h, isolated mRNA, and screened for ER^UPR^ markers. As supported by the cell proliferation data, glucose uptake inhibition yielded no appreciable activation of *ATF4* or *DDIT3* expression (Fig. S1D), which contrasted to a known inducer of ER^UPR^ (Tunicamycin) which activated ER^UPR^ and cell cycle gene signatures (Figs. S1E-G). Furthermore, increasing concentrations of KL-11743 did not induce apoptosis after 24 h (Figs. 1J–L), and the highest of these concentrations did not induce apoptosis after 48 or 72 h (Figs. 1M–O).

To confirm that a decrease in glucose uptake also reduced glycolysis, we conducted media analysis to quantify levels of glucose and lactate. As expected, treatment with KL-11743 abolished glucose consumption and decreased the concomitant production of lactate, the major metabolite formed by glycolysis in cancer cells [33](Fig. 2A). The glycolytic capacity of cells treated with KL-11743 was assessed using an Agilent Glycolysis Stress Test, where the extracellular acidification rate (ECAR) was measured as an indicator of the glycolytic rate. We demonstrated that KL-11743 ablated cellular response to glucose and oligomycin-stimulated glycolysis (Figs. 2B–D); whereas a structurally-related enantiomer (KL-12023) did not block ECAR (Figs. S1H–M). Additionally, we corroborated these data by directly determining if KL-11743 restricted access to extracellular glucose. When NB4, THP-1, and MOLT-4 are serum-deprived, glucose supplementation is essential to maintain viability; when both are withdrawn, cells undergo rapid apoptosis (Figs. S1N–P). As glucose uptake is necessary to preserve survival in this scenario, we tested if KL-11743 blocks glucose-dependent survival. To examine, cells were cultured in serum-free conditions for 24 h, followed by either glucose-withdrawal or glucose supplementation ± KL-11743 for an additional 24 h before cell death quantification. In the absence of serum, glucose withdrawal led to massive cell death that was preventable by titrating back glucose, while the addition of KL-11743 completely prevented glucose-dependent viability (Figs. S1N–P). Together, these data suggest the effects of KL-11743 are specific to effectively block glucose uptake in leukemia cell models.

**Figure 2.**
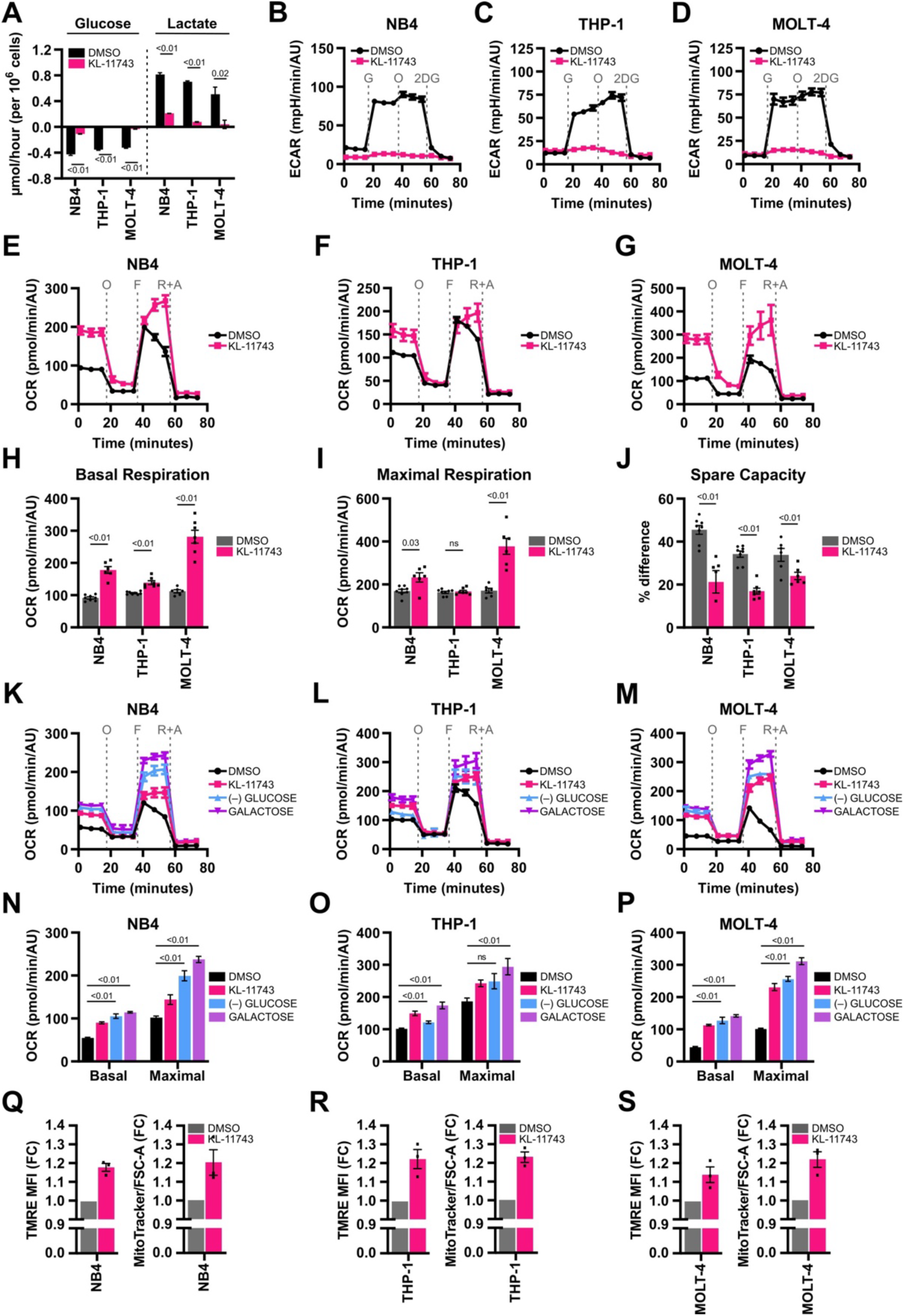
Inhibition of glucose uptake induces mitochondrial respiration as a compensatory bioenergetic mechanism. **(A)** NB4, THP-1, and MOLT-4 were treated with DMSO or KL-11743 (500 nM) for 24 h, followed by YSI media analysis of glucose and lactate concentrations. Average rates ± SEM, with negative values indicating consumption and positive values indicating secretion. *P* values were computed using an unpaired T test. **(B–D)** NB4, THP-1, and MOLT-4 were treated with DMSO or KL-11743 (500 nM) for 24 h and extracellular acidification rates (ECAR) were measured following an Agilent XF Glycolysis Stress Test. Injection of glucose (G, 10 mM) stimulated glycolysis, injection of oligomycin (O, 1 μM) stimulated maximal glycolytic rates, and injection of 2-deoxy-D-glucose (2DG, 50 mM) suppressed glycolysis. **(E–G)** Cells were treated as in *B*–*D* and OCR was measured using an Agilent XF Cell Mito Stress Test. **(H)** Quantification of basal respiration from *E*–*G*, calculated as the average of the 3 reads preceding injection of oligomycin (O). **(I)** Quantification of maximal respiration from *E*–*G*, calculated as the average of the 3 reads immediately following injection of FCCP (F). **(J)** Quantification of spare respiratory capacity from *E*–*G*, calculated as the percent difference between basal and maximal respiration reads. **(K–M)** NB4, THP-1, and MOLT-4 were treated with DMSO, KL-11743 (500 nM), cultured in glucose-deficient media ((−)GLUCOSE), or cultured in glucose-deficient media supplemented with 2 g/L galactose (GALACTOSE) for 24 h, followed by Agilent XF Cell Mito Stress Test. **(N–P)** Quantification of basal and maximal respiration levels from *K-M* graphed as in *H* and *I*. O: oligomycin (1 μM), F: FCCP (1 μM), R+A: rotenone (0.5 μM) + antimycin A (0.5 μM). Statistical significance for these data (panels *E-N*) was determined using non-parametric Mann-Whitney U test where *p* values are indicated. **(Q–S)** NB4, THP-1, and MOLT-4 were treated with DMSO or KL-11743 (500 nM) for 24 h, followed by flow cytometry analyses of mitochondrial membrane potential (TMRE) and mitochondrial mass (MitoTracker^TM^ Green). Data reported as the fold change (FC) to vehicle-treated samples. Panels *B*–*P* are presented as mean values ± SEM of at least 5 technical replicates; panels *Q*–*S* are mean values ± SEM of 3 technical replicates. Individual dots in bar graphs represent technical replicates.

Due to the dramatic decrease in glycolysis and continued growth of KL-11743-treated cells, we hypothesized that KL-11743 induces a metabolic reprogramming to maintain sufficient ATP pools through non-glycolytic pathways. The most efficient source of cellular ATP production is through mitochondrial respiration, so we evaluated mitochondrial function of KL-11743-treated cells using an Agilent XF Cell Mito Stress Test. Oxygen consumption rates (OCR) were measured as follows: after determining basal OCR, oligomycin (O) was used to inhibit ATP synthase, which depletes ATP-linked respiration; FCCP (F) was then used to transport hydrogen ions across the IMM down the concentration gradient, forcing maximal ETC activity; lastly, a combination of rotenone and antimycin-A (R+A) was used to inhibit Complex I (CI) and Complex III (CIII), respectively, to prevent *e^−^* shuttling thus revealing proton-leak and/or non-mitochondrial OCR (Figs. 2E–G). We found that KL-11743 significantly increased basal mitochondrial respiration in all cell lines (Fig. 2H). NB4 and MOLT-4 cells showed an increase in maximal respiration, indicating that treatment with KL-11743 enhances the maximal mitochondrial respiratory potential (Fig. 2I). We calculated the spare respiratory capacity as the % difference between basal and maximal respiration and observed a significant reduction in spare capacity in all cell lines, indicating that mitochondria of KL-11743-treated cells operate closer to their maximal respiratory potential (Fig. 2J).

We hypothesized that the decrease in glycolytic rate causes the increase in oxidative phosphorylation (OXPHOS). To test this, OCR was measured in cells cultured in glucose-free media (Figs. 2K–M). Basal OCR significantly increased in all cell lines when glucose was absent, compared to control conditions. Maximal respiration increased in glucose-free media in NB4 (Fig. 2N) and MOLT-4 (Fig. 2P), while failing to significantly change in THP-1 (Fig. 2O). THP-1 cells were the least glycolytic (Fig. 1A) and rely heavily on OXPHOS [34], leading them to be least affected by glucose depletion. We also compared mitochondrial respiration levels when we supplemented glucose-free media with galactose. Galactose is metabolized to glucose-6-phosphate through the Leloir pathway and enters glycolysis slower than glucose, preventing cells from using glycolysis as a rapid ATP source [35, 36]. As a result of decreased glycolytic rates, we expected cells to switch to increased mitochondrial respiration. We confirmed that across cell lines, glucose-free medium supplemented with galactose increased both basal and maximal OCR (Figs. 2N–P). These findings indicate that decreased glycolytic rates, either due to glucose transporter inhibition or substrate limitation, cause a compensatory increase in mitochondrial respiration. Finally, the positive effect of KL-11743 on mitochondrial respiration was also associated with increased mitochondrial delta psi, as measured by cellular TMRE staining, and increased mitochondrial mass (*i.e.,* MitoTracker^TM^ Green) (Figs. 2Q–S).

### The metabolic adaptation to KL-11743 requires a functional TCA cycle to fuel the electron transport chain

We next sought to define a metabolic signature for KL-11743 treated cells by performing targeted metabolomics primarily focused on carbohydrate metabolism. NB4 cells treated with KL-11743 demonstrated a marked loss in glycolytic intermediates yet with minimal effects on ATP levels (Figs. S2A–E). To identify conserved changes contributing to mitochondrial respiration in glucose uptake-inhibited cells, we compared metabolomic changes in NB4 and MOLT-4 cells, the two cell lines that showed the most compelling mitochondrial response to KL-11743. Among the enriched metabolite sets, alanine, aspartate, and glutamate metabolism ranked third-most enriched (Fig. S3A). Within this set, the metabolite showing the highest significant enrichment was aspartic acid (Fig. 3A). Aspartic acid plays a crucial role in the malate-aspartate shuttle, a pathway responsible for transporting *e^−^*from NADH produced in the cytosol (*i.e.*, by glycolysis) into mitochondria. It has been shown that KL-11743 synergizes with mutations in GOT1, a glutamate oxaloacetate transaminase used in the shuttle, in a colorectal carcinoma cell line [26]. To determine whether our model system displays a dependency upon glucose uptake inhibition, we used the malate-aspartate shuttle inhibitor 3-nitropropionic acid (3-NPA). 3-NPA impedes the TCA cycle, thereby preventing the formation of the required concentration of oxaloacetate to maintain the malate-aspartate shuttle flux [37]. We examined the mitochondrial respiratory function of co-treatment with 3-NPA and KL-11743 and observed that 3-NPA’s phenotype is dominant in NB4, decreasing basal and maximal OCR despite the presence of KL-11743 (Fig. 3B). This result led us to hypothesize that 3-NPA may be interfering with a pathway or enzyme that is upregulated in cells treated with KL-11743 and responsible for the increased mitochondrial oxidative phosphorylation. To investigate the potential bioenergetic dependency conferred by treatment with KL-11743 on the malate-aspartate shuttle, we quantified apoptosis following single and combined treatments of KL-11743 and 3-NPA. Our findings revealed no induction of apoptosis under these conditions (Fig. S3B).

**Figure 3.**
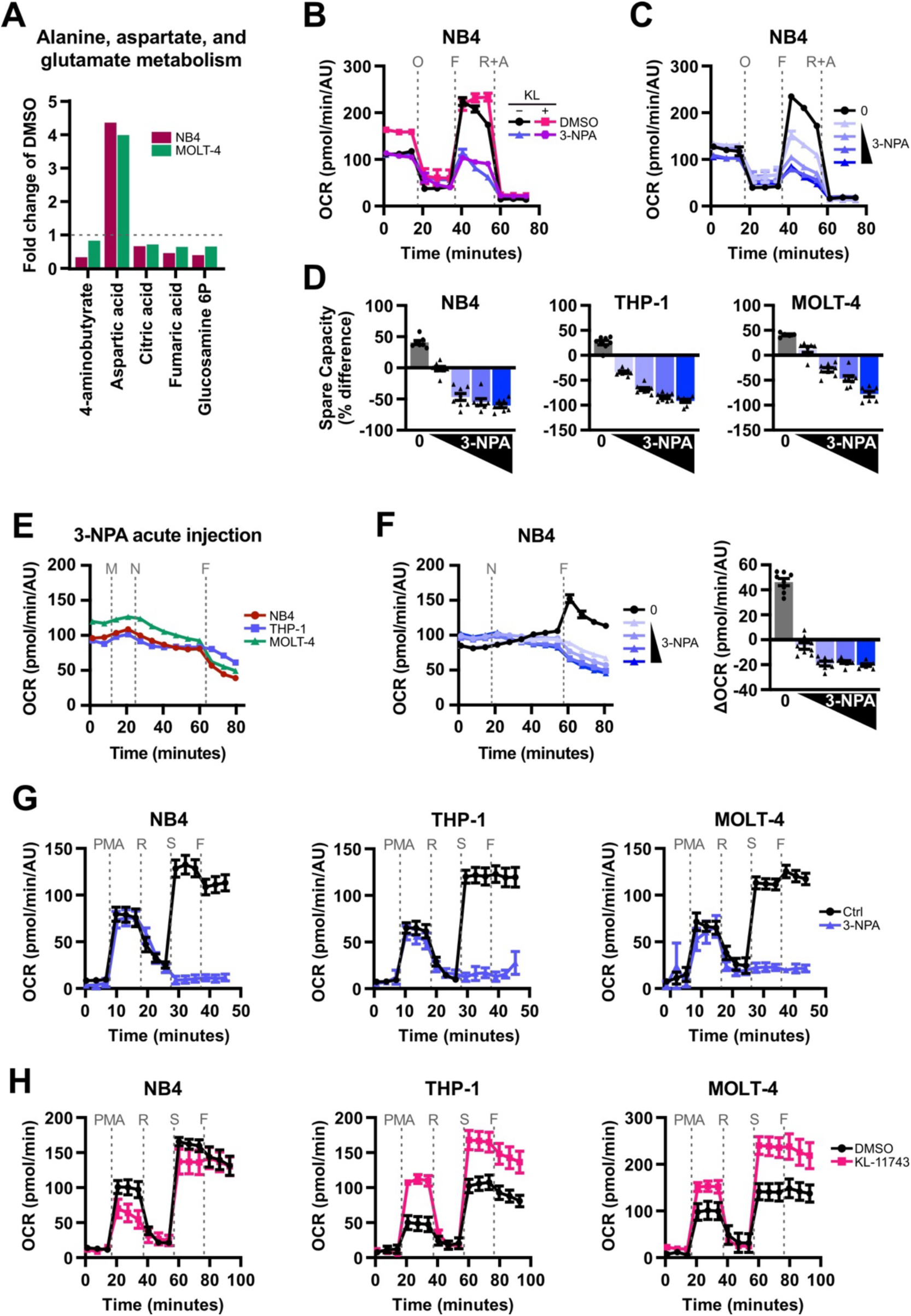
3-NPA prevents mitochondria from achieving maximal respiration and reverses the bioenergetic adaptation to KL-11743. **(A)** Metabolites that were significantly altered (*p*<0.05) in alanine, aspartate, and glutamate metabolism set in Figure S3A. **(B)** NB4 were treated with DMSO, KL-11743 (500 nM) ± 3-NPA (1 mM) for 24 h, and OCR was measured following an Agilent XF Cell Mito Stress Test. O: oligomycin (1 μM), F: FCCP (1 μM), R+A: rotenone (0.5 μM) + antimycin A (0.5 μM). **(C)** NB4 were treated with 3-NPA (10, 50, 100, 250 μM) for 24 h and OCR was measured. O: oligomycin (1 μM), F: FCCP (1 μM), R: rotenone (0.5 μM), A: antimycin A (0.5 μM). **(D)** NB4, THP-1, and MOLT-4 were treated with 3-NPA (10, 50, 100, 250 μM) for 24 h and OCR was measured following an Agilent XF Cell Mito Stress Test. Spare respiratory capacity calculated as the percent difference between basal and maximal OCR. **(E)** Changes in OCR were measured using an Agilent XFe96 Analyzer after injection of XF RPMI medium (M), 3-NPA (N, 1 mM), and FCCP (F, 1 μM). **(F)** Changes OCR of NB4 were measured using an Agilent XFe96 Analyzer after injection with 3-NPA (N, 250, 500, 750, 1000 μM) and FCCP (F, 1 μM). Right panel depicts the difference in OCR between the measurement immediately following FCCP injection and the measurement immediately preceding FCCP injection. **(G)** CI analysis of NB4, THP-1, and MOLT-4 treated with 3-NPA (1 mM) for 24 h. OCR was measured by an Agilent XFe96 Analyzer during sequential administration of a combination of PMA: pyruvate (10 mM) + malate (0.5 mM) + ADP (4 mM), R: rotenone (1 μM), S: succinic acid (10 mM), and F: FCCP (1 μM). **(H)** CI analysis of NB4, THP-1, and MOLT-4 treated with DMSO or KL-11743 (500 nM) for 24 h. OCR was measured by an Agilent XFe96 Analyzer during sequential administration of a PMA: pyruvate (10 mM) + malate (0.5 mM) + ADP (4 mM), R: rotenone (1 μM), S: succinic acid (10 mM), and F: FCCP (1 μM). Data are presented as the mean of 3 replicates ± SEM. Panels *B-H* are displayed as mean values of at least 6 technical replicates ± SEM.

To further assess potential combinatorial effects of KL-11743 and 3-NPA, we employed Single-Cell and Population-Level Analyses Using Real-Time Kinetic Labeling (SPARKL)[29], which allowed us to assess both cell death kinetics and morphology. We again saw no induction of cell death when 3-NPA treatment was combined with KL-11743 (Fig. S3C). However, combined treatment prevented the formation of cell clumps in culture, indicating stress and/or decreased proliferation (Fig. S3D). We also evaluated two other inhibitors that target the malate-aspartate shuttle through distinct mechanisms: phenylsuccinic acid (PSA) and aminooxyacetic acid (AOAA). PSA functions by blocking the mitochondrial malate-α-ketoglutarate carrier. AOAA, a carbonyl-trapping agent, inhibits enzymes dependent on pyridoxal phosphate (PLP), including the aspartate aminotransferase employed in the malate-aspartate shuttle [37]. None of the malate-aspartate shuttle inhibitors induced cell death when combined with KL-11743, demonstrating that KL-11743 did not induce a dependency on the malate-aspartate shuttle (Figs. S3E-F).

Subsequently, our focus shifted towards the mitochondrial bioenergetics phenotype following treatment of 3-NPA (Fig. 3B). As an inhibitor of succinate dehydrogenase, 3-NPA completely abolished CII-dependent OCR, measured following succinate and ADP addition in the presence of rotenone (Fig. S4A). To determine the effects of 3-NPA on overall mitochondrial respiration, we treated NB4 cells with a titration of 3-NPA for 24 h and observed a dose-dependent inhibition of FCCP-induced OCR (Fig. 3C). The same experiment was also performed comparing NB4, MOLT-4, and THP-1, and while basal respiration levels were largely unaltered, maximal OCR markedly decreased and resulted in negative spare capacity values for all cell lines (Fig. 3D). We showed that this response is not an adaptation but rather an acute response to succinate dehydrogenase inhibition, as FCCP injection ∼40 min after 3-NPA injection also failed to increase OCR (Fig. 3E). A low-dose titration of 3-NPA in NB4 revealed an acute, dose-dependent inhibition of FCCP-induced maximal OCR, whereas untreated cells responded to FCCP as expected (Fig. 3F). Notably, this inhibition of FCCP-induced maximal OCR was not observed in treatment with TTFA, an inhibitor of *e^−^* transfer within CII but not of the enzyme succinate dehydrogenase, which also fails to induce apoptosis when combined with KL-11743 (Fig. S4B). Prolonged TTFA treatment dramatically decreased basal OCR, but cells were able to increase OCR following FCCP treatment (Fig. S4C-D). These results indicate that the observed phenotype of 3-NPA is a consequence of inhibiting a TCA cycle enzyme, not of directly inhibiting or altering functionality of the ETC. We next measured Complex-specific oxygen consumption rates, where we permeabilized the plasma membrane and directly supplied substrates for oxidation. We observed that when substrates are provided to 3-NPA treated cells, energetics are remarkably similar to controls, indicating that 3-NPA is acting to limit the ETC substrates required to reach maximal respiration rates (Fig. 3G). To assess how CI may be functioning in 3-NPA-treated cells in culture, we assessed whole-cell [NAD^+^] and [NADH]. We found that NAD^+^/NADH ratios increased after 3-NPA treatment, indicative of a change in redox flux and possibly increased NADH oxidation by CI of the ETC (Fig. S4E). Interestingly, treatment with KL-11743 showed similar trends (Figs. S4E–F). We performed a CI-specific OCR assay on cells treated with KL-11743 for 24 h. THP-1 and MOLT-4 cells exhibited increased CI activity, while NB4 cells were minorly affected. KL-11743-treated THP-1 and MOLT-4 cells also displayed increased CII activity (depicted as OCR levels following succinate injection), indicating an overall enhancement in mitochondrial ETC activity and/or efficiency driven in part by an enhanced substrate availability (Fig. 3H).

### KL-11743-mediated induction of mitochondrial oxygen consumption requires glutamine

Glutamine is a critical substrate that drives TCA cycle flux, with many leukemias displaying a dependency and metabolic addiction [38, 39]. We observed that KL-11743 treatment increased glutamate export, suggesting that KL-11743 may induce an intracellular re-routing of glutamine metabolism (Figs. 4A, S5A). Therefore, we hypothesized that the metabolic reprogramming induced by KL-11743 may be glutamine dependent. To initially investigate this potential interaction, we examined the cellular consequences of glutamine deprivation (-Q) ± KL-11743 treatment. After 24 hours, we observed a marked reduction in cellular viability with glutamine deprivation alone, which could only be minorly influenced by KL-11743 addition (Fig. 4B). These results demonstrated that glutamine deprivation studies were limited to only a few hours to avoid confounding influences of cell death.

**Figure 4.**
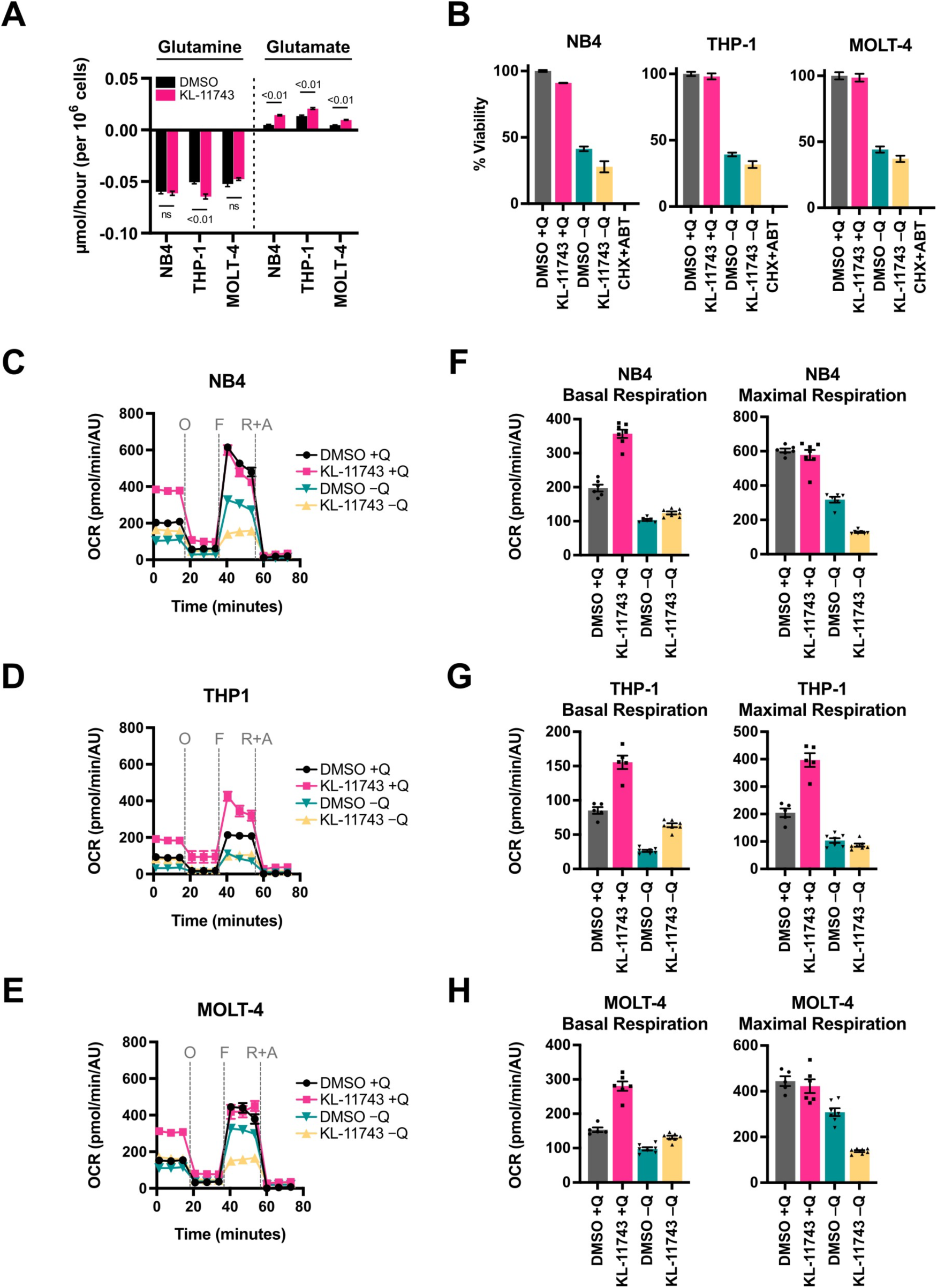
Glutamine metabolism is altered in KL-11743 treated cells. **(A)** NB4, THP-1, and MOLT-4 were treated with DMSO or KL-11743 (500 nM) for 24 h, followed by YSI media analysis of glutamine and glutamate concentrations. Average rates are graphed ± SEM, with negative values indicating consumption and positive values indicating secretion. **(B)** NB4, THP-1, and MOLT-4 were cultured in complete media ((+)Q) or glutamine-free ((−)Q) media treated with DMSO or KL-11743 (500 nM) for 24 h. Viability was determined by CellTiter Glo kit. **(C-E)** NB4, THP-1, and MOLT-4 were treated with DMSO or KL-11743 (500nM) for 24 h prior to changing them into glutamine-free ((−)Q) media for 6 h. OCR was measured using an Agilent XF Cell Mito Stress Test. O: oligomycin (1 μM), F: FCCP (1 μM), R+A: rotenone (0.5 μM) + antimycin A (0.5 μM). **(F)** Quantification of basal and maximal respiration from *C*, calculated as the average of the 3 reads preceding injection of oligomycin (O), and as the average of the 3 reads immediately following infection of FCCP (F), respectively. **(G)** Quantification of basal and maximal respiration from *D*, calculated as the average of the 3 reads preceding injection of oligomycin (O), and as the average of the 3 reads immediately following infection of FCCP (F), respectively. **(H)** Quantification of basal and maximal respiration from *E*, calculated as the average of the 3 reads preceding injection of oligomycin (O), and as the average of the 3 reads immediately following infection of FCCP (F), respectively.

To determine if the KL-11743 increased OCR and altered bioenergetics were dependent on glutamine (Figs. 2E-P), we treated cells with KL-11743 for 18 hours before removing glutamine for the final 6 hours (*n.b.,* cells are 100% viable after 6 hours without glutamine, data not shown), and then measured OCR. As expected, we observed that KL-11743 treatment markedly elevated basal respiration rates in the presence of glutamine; yet in the absence of glutamine, KL-11743-stimulated OCR was severely blunted and led to a reduction in maximal respiratory capacity (Figs. 4C-H). Additionally, acute glutamine withdrawal for only 2 hours prior to the Agilent assay was sufficient to erode KL-11743-stimulated OCR (Fig. S5B). These findings led us to conclude that KL-11743-mediated induction of mitochondrial oxygen consumption requires glutamine.

### Inhibition of glucose uptake results in a reliance on Complex I function for survival

All substrates that drive TCA flux and NADH regeneration converge on Complex I (CI) of the ETC. To investigate KL-11743-induced changes in CI activity, we used an in-gel activity assay to evaluate endogenous CI. NB4, THP-1, and MOLT-4 were treated with KL-11743 for 24 h, and the resulting heavy membrane fractions were extracted for intact CI activities. We observed an increase in CI activity in THP-1 and MOLT-4 upon KL-11743 treatment (Fig. 5A; arrow indicates ASSEMBLED). Additionally, we detected activity below the arrow, which is defined as “FREE” — not fully-assembled CI subunits from the *N* module, still capable of oxidizing NADH [40](and data not shown). Quantification of the free versus assembled bands revealed that KL-11743-treated THP-1 and MOLT-4 cells increased CI expression and assembly, while NB4 cells exhibited decreased activity of free CI (Fig. 5B). KL-11743 treatment also led to increased transcription of mitochondrially-encoded genes, including those encoding Complex I (Fig. 5C). These findings corroborate CI-dependent oxygen consumption rates observed following KL-11743 treatment (Fig. 3H).

**Figure 5.**
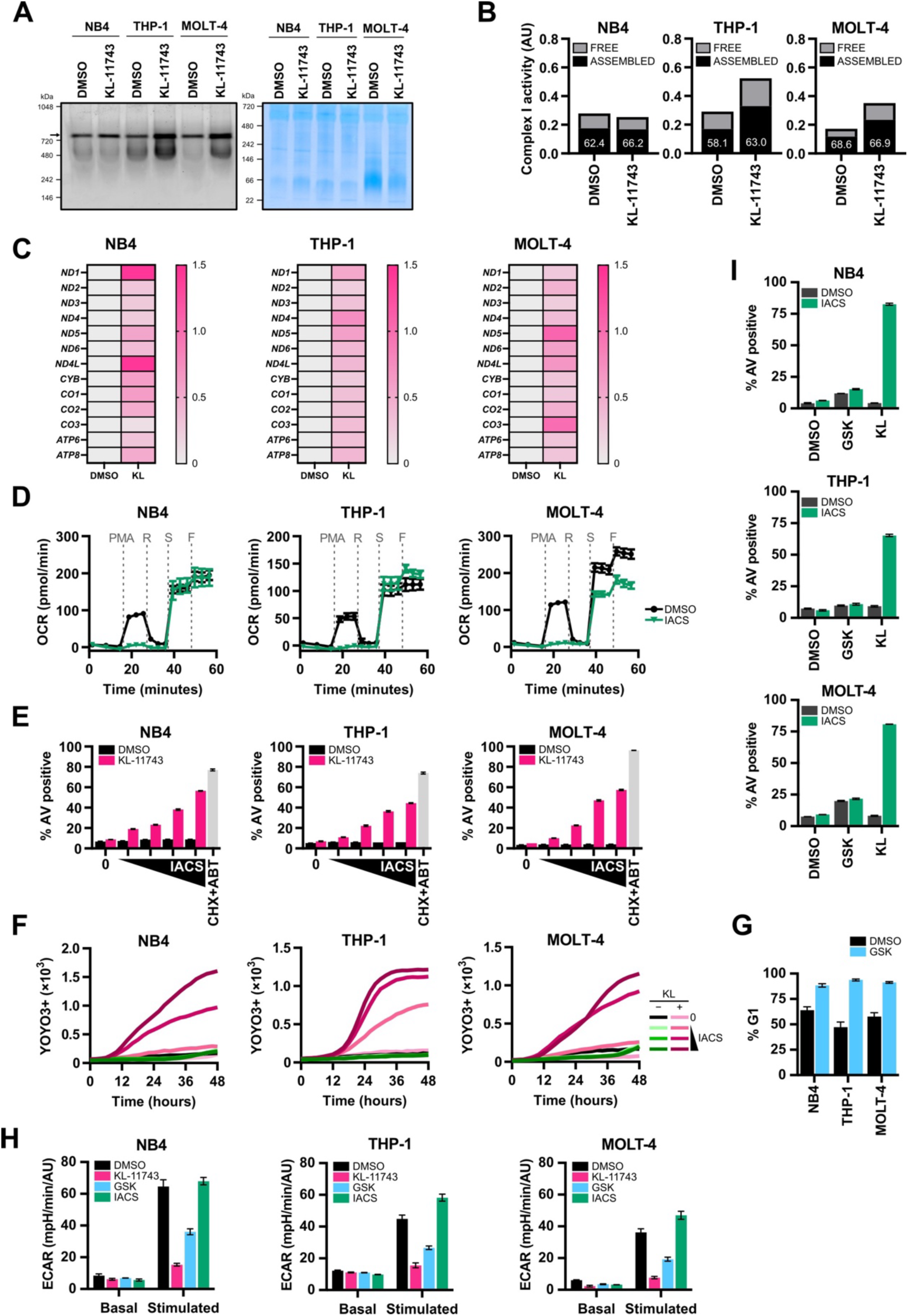
Complete inhibition of glucose uptake promotes Complex I dependency for survival. **(A)** NB4, THP-1, and MOLT-4 were treated with DMSO or KL-11743 (500 nM) for 24 h. Heavy membranes were isolated, digitonin-extracted, and the protein complexes were resolved using native PAGE conditions. In-gel activity was stimulated with NADH and the arrow above 720 kDa indicates assembled CI; the right panel is a Coomassie Blue loading control. **(B)** Quantification of the main bands indicated by the arrow (ASSEMBLED) and the activity detected below the assembled bands (FREE) in *A*. Numbers depicted in bars show the % of total activity attributed to the assembled CI species. **(C)** NB4, THP-1, and MOLT-4 were treated with DMSO or KL-11743 (500 nM) for 24 h, and total RNA was harvested. The fold change of transcripts from mitochondria encoded genes were measured by real-time qPCR. Expression was normalized against *18S*. **(D)** CI analysis of NB4, THP-1, and MOLT-4 treated with DMSO or IACS (10 nM) for 24 h as in *C*–*E*. **(E)** NB4, THP-1, and MOLT-4 were treated with DMSO, KL-11743 (500 nM), or IACS (1, 3, 6, 10 nM) ± KL-11743 (500 nM) for 24 h. Apoptosis was measured by AV labeling and flow cytometry. CHX (50 μg/mL) + ABT-737 (1 μM) is a positive control for apoptosis. **(F)** NB4, THP-1, and MOLT-4 were treated as with DMSO, KL-11743 (500 nM), or IACS (1, 3, 6, 10 nM) ± KL-11743 (500 nM), imaged every 2 h, and analyzed for YOYO3+ cells; the mean YOYO3+ events per image of 2 replicates is presented. **(G)** NB4, THP-1, and MOLT4 were treated with DMSO or GSK-1120212 (25 nM) for 24 h before PI staining and flow cytometry. **(H)** NB4, THP-1, and MOLT4 were treated with GSK-1120212 (25 nM), KL-11743 (500 nM), or IACS (10 nM) for 24 h, and ECAR (basal versus glucose-stimulated, 10 mM) was measured. **(I)** NB4, THP-1, and MOLT4 cells were treated with GSK-1120212 (25 nM) ± IACS (10 nM) for 24 h. Apoptosis was measured by AV labeling and flow cytometry. Error bars are the average of 3–6 technical replicates ± SEM.

Considering the increased expression, assembly, and/or function of CI in most of the KL-11743-treated models, we investigated whether this phenomenon indicated a metabolic dependency on CI. To test this, we treated cells for 24 h with IACS-010769 (IACS), a clinically relevant CI inhibitor [41], and then performed OCR and cell death studies. A CI-specific OCR assay revealed that IACS markedly disrupts CI function without inhibiting CII (Fig. 5D). Also, the combination of KL-11743 with IACS resulted in a dose- and time-dependent synthetic lethality, as demonstrated by Annexin V and cell death kinetics studies, respectively, revealing a potent requirement for CI function when cells are exposed to KL-11743 (Figs. 5E-F). As a comparison, small molecules (*e.g.,* GSK-1120212 / Trametinib) that inhibit oncogenic MAPK signaling also block glucose uptake, but through the reduction of GLUTs on the cell surface. NB4, THP1, and MOLT4 harbor activated RAS, and 24 h exposure to GSK-1120212 (GSK) caused G1 accumulation, which demonstrates a potent drug response (Fig. 5G). The same treatment also reduced glucose-stimulated ECAR by ∼50%, yet KL-11743 treatment eliminated ECAR by ∼90% (Fig. 5H). To determine if GSK-mediated loss of ECAR also results in CI dependency for survival, cells were co-treated with GSK and IACS for 24 h, then stained for AV and flow cytometry. Despite a reduction in glycolysis, GSK treatment did not synergize with IACS to promote cell death (Fig. 5I) suggesting that either a partial loss of ECAR is insufficient to create dependency or direct GLUT inhibition is required for optimal cell death responses.

### AML patient-derived cells demonstrate potential clinical utility of combined KL-11743 treatment with Complex I inhibition

To explore a potential pre-clinical implication of glucose uptake inhibition, we investigated its effects in patient-derived cells. First, we utilized induced pluripotent stem cell (iPSC) lines derived from normal cells, as well as from a patient with AML, including isogenic lines with and without a KRAS^G12D^ mutation (AML-4.24: KRAS^WT^ AML and AML-4.10: KRAS^G12D^ AML) [27, 28]. We analyzed RNA-seq data from CD34+ normal hematopoietic stem/progenitor cells (HSPCs) or AML leukemia stem cells (LSCs) derived through *in vitro* differentiation from these lines (GSE92494; Fig. 6A). We noted that several mitochondria-related pathways are significantly downregulated in AML LSCs, compared to normal HSPCs, including TCA, OXPHOS, mitochondrial protein import, and mitochondrial protein translation (Figs. 6B– E). We also noted that glycolysis genes and *SLC2A1* (GLUT1, a KL-11743 target), are upregulated (Fig. 6F), highlighting both contributions and therapeutic potential of the pathway.

**Figure 6.**
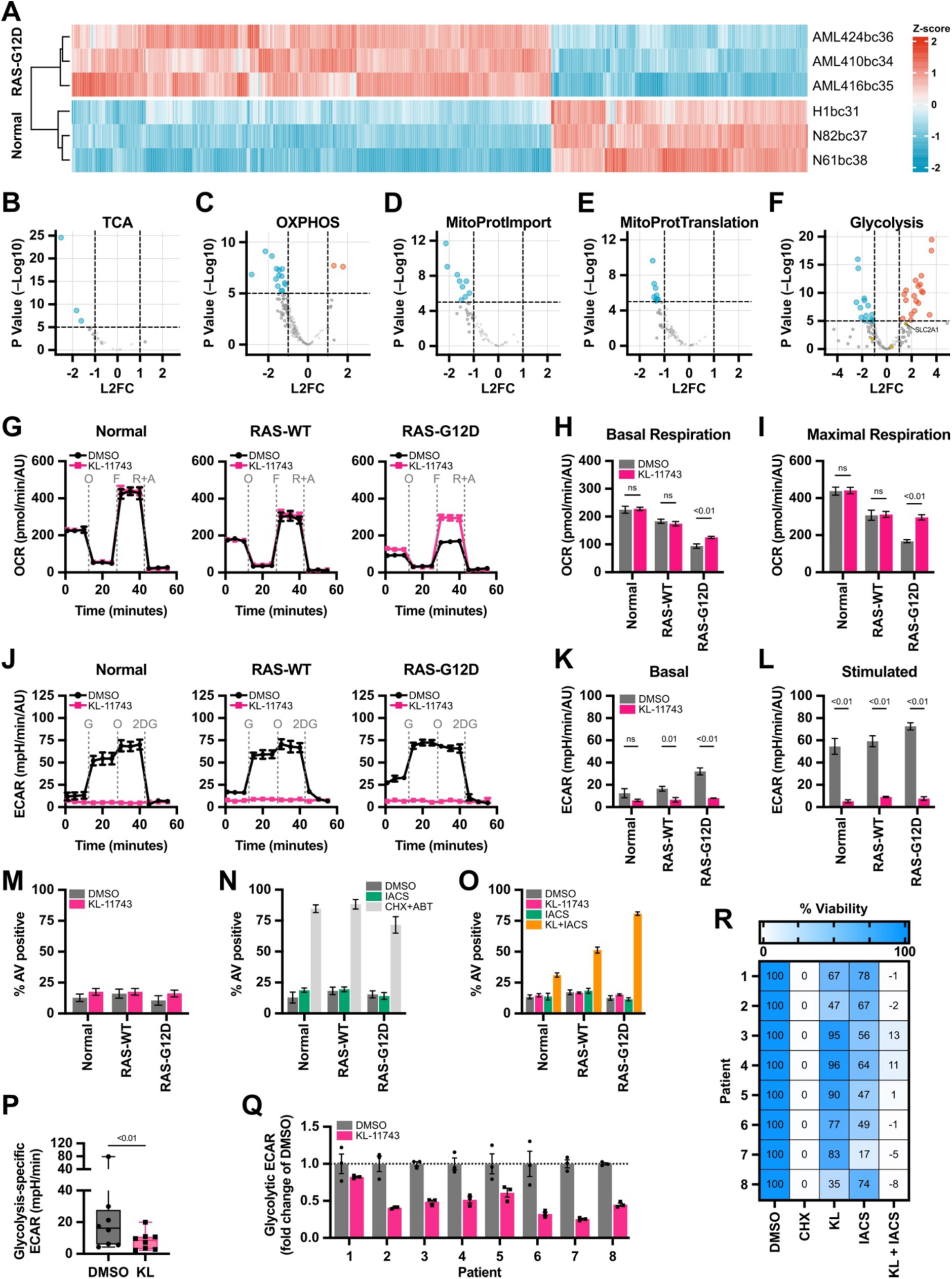
Patient-derived cells demonstrate clinical utility of combined KL-11743 treatment with Complex I inhibition. **(A)** Differential gene expression analysis comparing normal HSPCs, derived from the human embryonic stem cell (hESC) line H1 and two iPSC lines (N-8.2 and N-6.1); KRAS^WT^ AML-iPSC-derived LSCs (AML-4.24 and AML-4.16) and KRAS^G12D^ AML-iPSC derived LSCs (AML-4.10). Heatmap showing expression of 1447 genes differentially expressed between normal iPSC-HSPCs and AML-iPSC-LSCs using scaled log normalized counts. **(B–F)** RNA-seq data from *A* was re-analyzed to directly compare normal iPSC-HSPCs to AML-iPSC LSCs. Volcano plots represent the log2 fold-change (FC) and *p*-value of genes enriched in the indicated gene ontology pathways. *SLC2A1*, a KL-11743 target, is highlighted. The number of genes compared in each pathway as follows: TCA, 28; OXPHOS, 199; Mitochondrial Protein Import: 61; Mitochondrial Protein Translation: 94; Glycolysis: 193. **(G)** Normal HSPCs or AML LSCs derived from iPSC lines N-8.2, AML-4.24 (KRAS^WT^) and AML-4.10 (KRAS^G12D^) were treated with DMSO or KL-11743 (500 nM) and OCR was measured following an Agilent XF Cell Mito Stress Test. O: oligomycin (1 μM), F: FCCP (1 μM), R+A: rotenone (0.5 μM) + antimycin A (0.5 μM). **(H)** Quantification of basal respiration from *G*, calculated as the average of the 3 reads preceding injection of oligomycin (O). **(I)** Quantification of maximal respiration from *G*, calculated as the average of the 3 reads immediately following injection of FCCP (F). **(J)** Normal HSPCs or AML LSCs derived from iPSC lines N-8.2, AML-4.24 (KRAS^WT^) and AML-4.10 (KRAS^G12D^) were treated with DMSO or KL-11743 (500 nM) for 24 h and ECAR was measured following an Agilent XF Glycolysis Stress Test. G: glucose (10 mM), O: oligomycin (1 μM), 2DG: 2-deoxy-D-glucose (50 mM). **(K–L)** Quantification of basal and glucose-stimulated ECAR from *H*, calculated as the average of the 3 reads immediately before and following injection of glucose (G), respectively. **(M)** Normal HSPCs or AML LSCs derived from iPSC lines N-8.2, AML-4.24 (KRAS^WT^) and AML-4.10 (KRAS^G12D^) were treated with DMSO or KL-11743 (500 nM) for 24 h. Apoptosis was measured by AV labeling and flow cytometry. Data are presented as the mean of 3 replicates ± SD. **(N)** Cells were treated and analyzed as in *M*, but with IACS (10 nM). CHX (50 μg/mL) + ABT (1 μM) is a positive control for apoptosis. Data are presented as the mean of 3 replicates ± SD. **(O)** Cells were treated and analyzed as in *M-N*, but with KL-11743 (1 μM) ± IACS (10 nM). Data are presented as the mean of 3 replicates ± SD. **(P)** Primary cells from AML patients 1–8 were treated with DMSO (0.1%) or KL-11743 (500 nM) at a density of 100,000 cells/mL, cultured for 24 h, and ECAR was measured following an Agilent XF Glycolysis Stress Test. Data are presented as the glycolysis-specific ECAR, calculated as the difference between basal and glucose-stimulated ECAR using the timepoint immediately before and after the glucose injection. **(Q)** Data from *P* presented as the fold change relative to DMSO. Data are the mean of 3 replicates ± SEM. **(R)** Primary cells from AML patients 1–8 were treated with DMSO (0.1%), KL-11743 (500 nM), IACS (10 nM), CHX (50 μg/mL), or indicated combination for 24 h before viability was measured. DMSO and CHX are the negative and positive cell death controls. Data were normalized to the DMSO and CHX treatments as 100 and 0%, respectively.

We next compared the effects of KL-11743 treatment on mitochondrial bioenergetics in normal HSPCs and AML LSCs using the Agilent XF Cell Mito Stress Test. We observed that basal OCR was highest in normal HSPCs and lowest in KRAS^G12D^ LSCs. In normal HSPCs and KRAS^WT^ AML LSCs, KL-11743 treatment did not influence basal or maximal OCR (Figs. 6G-I). In contrast, KRAS^G12D^ LSCs demonstrated a marked induction of OCR following KL-11743 treatment suggesting oncogenic MAPK signaling suppresses OCR, but KL-11743 treated KRAS^G12D^ cells respond by inducing mitochondrial bioenergetics (Figs. 6G-I), similarly to our observations in previous cellular models (Fig. 2). In the iPSC-derived cells, OCR was also inversely proportional to ECAR with KRAS^G12D^ LSCs demonstrating the highest rates of glycolysis, and independent of genotype, KL-11743 treatment completely inhibited basal- and glucose-stimulated ECAR (Figs. 6J–L). Akin to previous data (Figs. 1J–O, 5I), KL-11743 or IACS single treatments failed to induce apoptosis in the iPSC-derived HSPCs/LSCs (Figs. 6M–N). Given the KL-11743 specific induction of OCR in KRAS^G12D^ cells, we hypothesized that loss of glucose uptake would create the strongest dependency for mitochondrial respiration for survival in this genotype. Indeed, the combined treatment of KL-11743 and IACS induced marked apoptosis in the majority (>80%) of KRAS^G12D^ AML cells within 24 h, and while ∼50% of KRAS^WT^ AML cells also responded suggesting potential application to broader leukemia genotypes, the majority of normal HSPCs survived treatment (Fig. 6O).

To corroborate and extend the above observations, we treated primary cells from a diverse cohort of AML patient samples (Table 1, Supplementary Data File 1) with KL-11743 for 24 h and measured basal- and glucose-stimulated ECAR. Indeed, all patients responded to KL-11743 by reducing their basal and glucose-stimulated ECAR (Figs. 6P– Q, S6A–B). Next, we determined if the KL-11743 mediated reduction in ECAR created a metabolic vulnerability and sensitization to IACS. To do so, AML patient cells were treated with KL-11743 (500 nM) ± IACS (10 nM) for 24 h, and then analyzed by MTT for viability. KL-11743 treatments did impact on viability to different degrees in each patient; for example, patients 3, 4, 5, 6, and 7 displayed minimal viability changes after KL-11743 treatment, while patients 1, 2, and 8 exhibited ∼25–50% decreased viability (Figs. 6R, S6C). However, all patient samples responded similarly to KL-11743+IACS treatment, as this combination caused ∼90–100% death, suggesting the therapeutic potential of the pathway and vulnerability created by KL-11743 in AML (Figs. 6R, S6C).

## DISCUSSION

The results of our study demonstrate that KL-11743 effectively inhibits glucose uptake in a variety of acute leukemic cell lines, resulting in a reduction in glycolysis. Despite the highly glycolytic nature of these cells, treatment with KL-11743 did not induce apoptosis, indicating that glucose uptake inhibition alone is not sufficient to trigger cell death. To gain a deeper understanding of the metabolic consequences of glucose uptake inhibition, we investigated mitochondrial bioenergetics, targeted metabolic profiling, and nonbiased metabolomics. We showed that KL-11743 enhanced mitochondrial respiration in three highly glycolytic cell lines and we suggest that cells dependent on glycolysis rewire their metabolism in response to limited glucose, compensating for the loss of ATP.

Previous studies demonstrate that when cells are cultured in glucose-depleted medium, they exhibit increased formation of super-complexes composed of CI, CIII, and CIV, and favor CI substrate oxidation [31]. Our work supports the notion that CI substrate oxidation is a metabolic dependency when glucose uptake is inhibited, as KL-11743 increased the cytosolic NAD^+^/NADH ratio and inhibiting CI with IACS-010759 induced a synthetic lethality. Interestingly, the synthetic lethality with combined GLUT and CI inhibition was observed among all cell lines tested, regardless of whether their response to KL-11743 induced CI assembly and activity (THP-1 and MOLT-4), or induced OCR without substantially increased CI assembly and activity (NB4). When GLUT1-4 and CI are acutely inhibited, cells are bioenergetically stressed and commit to apoptosis.

With the development and initial characterization of KL-11743, Olszewski *et al*. (2022) suggested therapeutic potential for use of glucose uptake inhibition with *SDHB*-mutant tumors for a bioenergetic lethality and demonstrated decreased proliferation with KL-11743 treatment in these tumor types. Contrary to our expectations, we observed that inhibition of CII did not lead to cell death when combined with KL-11743, indicating limited therapeutic efficacy in *SDHB*-mutant tumors. While our models did not die upon combined GLUT1–4 and succinate dehydrogenase inhibition, cells were visibly stressed and could not reach maximal respiration rates. We propose that upon 3-NPA inhibition, the TCA cycle rearranges for continued synthesis of CI substrates. Literature supports that the TCA cycle can reroute during differentiation, bypassing succinate dehydrogenase by shuttling citrate out of mitochondria, where it can be converted oxaloacetate. Oxaloacetate is then converted into malate, which enters mitochondria and supplies the truncated TCA cycle [42]. Additionally, the TCA substrate α-ketoglutarate can be converted to oxaloacetate by GOT1. Oxaloacetate can be converted to citrate and contribute to a truncated TCA cycle, bypassing succinate dehydrogenase, or it can supply the malate-aspartate shuttle to bring NADH into mitochondria [37]. Our data show increased aspartate levels after KL-11743 treatment, suggesting increased malate-aspartate shuttle flux. Further investigations using ^13^C tracing to reveal the TCA metabolites generated after CII inhibition could determine whether succinate dehydrogenase inhibition allows AML cells to reroute their TCA cycles to produce NADH but not FADH_2_.

Previous studies have shown that AML cells possess lower spare respiratory capacities compared to hematopoietic cells, rendering them sensitive to increased oxidative metabolic stress, such as the induction of the TCA cycle and ETC through exogenous palmitate [43]. A low spare capacity may imply inefficiency, either in ETC arrangement or within the complexes themselves. When overstimulated or driven to work rapidly, the inefficient ETC may result in *e*^−^ leakage and ROS production. Across our cell lines, spare respiratory capacity was decreased in response to KL-11743, suggesting increased sensitivity to oxidative stress. It has been shown that mitochondrial ROS is increased by glucose limitation, but unaffected by IACS [44]. It would be interesting to investigate whether this is another vulnerability in KL-11743-treated cells, and whether overwhelming the ETC might synergize with the induction of oxidative stress and result in apoptosis. We have observed that cells cultured in substrate-limited growth media containing low concentrations of glucose, glutamine, and serum, and supplemented with L-carnitine, are sensitive to KL-11743 treatment (data not shown). Carnitine promotes fatty acid transport into mitochondria, and may be serving to overwhelm the ETC, accumulate ROS, and trigger cell death.

Our observations in acute leukemic cell lines expand upon recent investigations into BAY-876, a GLUT1 inhibitor. Rodriguez-Zabala *et al*. demonstrate that GLUT1 knockout decreases leukemia cell growth and induces apoptosis *in vivo*, and that combined BAY-876 and IACS treatment increases survival of tumorigenic mice, as well as viability of AML patient samples [45]. Nevertheless, a subset of patients were not responsive to combined GLUT1 and CI inhibition, potentially attributable to variances in GLUT expression profiles or transcriptional adaptations induced by GLUT1 inhibition [46, 47]

We propose that targeting this metabolic signature with KL-11743 holds promise as a therapeutic approach to treatment of hematological malignancies. Indeed, when we treated multiple models of leukemia ranging from standard laboratory cell lines, iPSC-based clones, and primary cells derived from AML patients, KL-11743 mediated inhibition of glucose uptake resulted in dependency upon CI for survival. IACS-010759, the CI inhibitor used in this study, has been reported to inhibit tumor growth in leukemia mouse models [41, 48, 49]. However, IACS-010759 encountered setbacks in phase 1 clinical trials due to dose-limiting toxicities impacting non-cancer cells, underscoring the necessity for comprehensive characterization of the systemic effects of metabolic inhibitors before advancing to clinical trials [50, 51]. Overall, our study begins to appreciate the metabolic consequences and vulnerabilities of KL-11743 treatment in cancer models. These findings provide insights into the metabolic adaptations and dependencies that arise in leukemic cells under glucose-deprived conditions, presenting potential therapeutic opportunities for clinical interventions.

## CONCLUSIONS

In conclusion, using a combination of targeted and unbiased metabolomics, real-time bioenergetics analyses, and high-content live-cell imaging in the context of multiple leukemic models, we methodically dissected the cellular responses to acute glucose uptake inhibition, all of which converge upon a glutamine-dependent bioenergetic vulnerability centered on mitochondrial Complex I. We demonstrated that oncogenic RAS expressing iPSC AML cells have the strongest dependency on Complex I for survival upon acute glucose uptake inhibition. Furthermore, we expanded the impact of this study by examining primary AML cells isolated from a cohort of Mount Sinai Health System patients to demonstrate the translational potential of dual inhibition of glucose uptake and Complex I, highlighting the relevance of these biological observations.

## ABBREVIATIONS

AML: acute myeloid leukemia
GLUT: glucose transporters
iPSC: induced pluripotent stem cell
NADH: nicotinamide adenine dinucleotide
TCA: tricarboxylic acid
ATP: adenosine triphosphate
FADH_2_: flavin adenine dinucleotide
FCCP: carbonyl cyanide-p-trifluoromethoxyphenylhydrazone
DMSO: dimethyl sulfoxide
SSC-A: side scatter-area
FSC-A: forward scatter-area
ER: endoplasmic reticulum
ECAR: extracellular acidification rate
OCR: oxygen consumption rates
O: oligomycin
R+A: rotenone + antimycin-A
OXPHOS: oxidative phosphorylation
3-NPA: 3-nitropropionic acid
SPARKL: single-cell and population-level analyses using real-time kinetic labeling
PSA: phenylsuccinic acid
AOAA: aminooxyacetic acid
PLP: pyridoxal phosphate
Q: glutamine
CI: Complex I
CII: Complex II
HSPCs: hematopoietic stem/progenitor cells
LSCs: leukemia stem cells

## DECLARATIONS

### Ethics Approval And Consent To Participate

De-identified patient samples were provided by the Tisch Cancer Institute’s Hematological Malignancies Tissue Bank through an Institutional Review Board at the Mount Sinai School of Medicine approved protocol (STUDY-11-02054-MOD008; PI: Bridget Marcellino).

### Consent For Publication

“Not applicable.”

### Data and Materials Availability

All data needed to evaluate the conclusions in the paper are present in the paper or the Supplementary Materials. The datasets used and/or analyzed during the current study are available from the corresponding author on reasonable request.

### Author Contributions

Conceptualization: MK, JK, JDG, APT, MVP, and JEC. Methodology: AGK and EPP. Validation: MK, JK, JDG, APT, AGK, and AB. Formal Analysis: MK, JK, JDG, JH, and AB. Investigation: MK, JK, JDG, APT, IA-E, JH, AE, AGK, CC, JA, SEG-B, AB, and YC. Resources: BKM, EPP, and MVP. Writing – Original Draft: MK and JEC. Writing – Review & Editing: MK, JK, JDG, and JEC. Visualization: MK and JDG. Supervision: JEC. Funding Acquisition: EPP and JEC.

## Acknowledgments

We thank the TCI Shared Resources (Flow Cytometry), the TCI Hematological Malignancies Tissue Bank at the Icahn School of Medicine at Mount Sinai, and the Department of Oncological Sciences for research support. Additionally, we wish to thank Metware Biotechnology, Dr. Irwin Kurland and the Stable Isotope & Metabolomics Core at the Albert Einstein College of Medicine, and the Donald B. and Catherine C. Marron Cancer Metabolism Center at Memorial Sloan Kettering Cancer Center for their assistance. A portion of this manuscript was adopted from Monika Komza’s Master’s Thesis, entitled “Inhibition of glucose uptake reveals adaptive metabolic reprogramming and a reliance on mitochondrial respiration.”

---

**Supplemental Figure 1.**
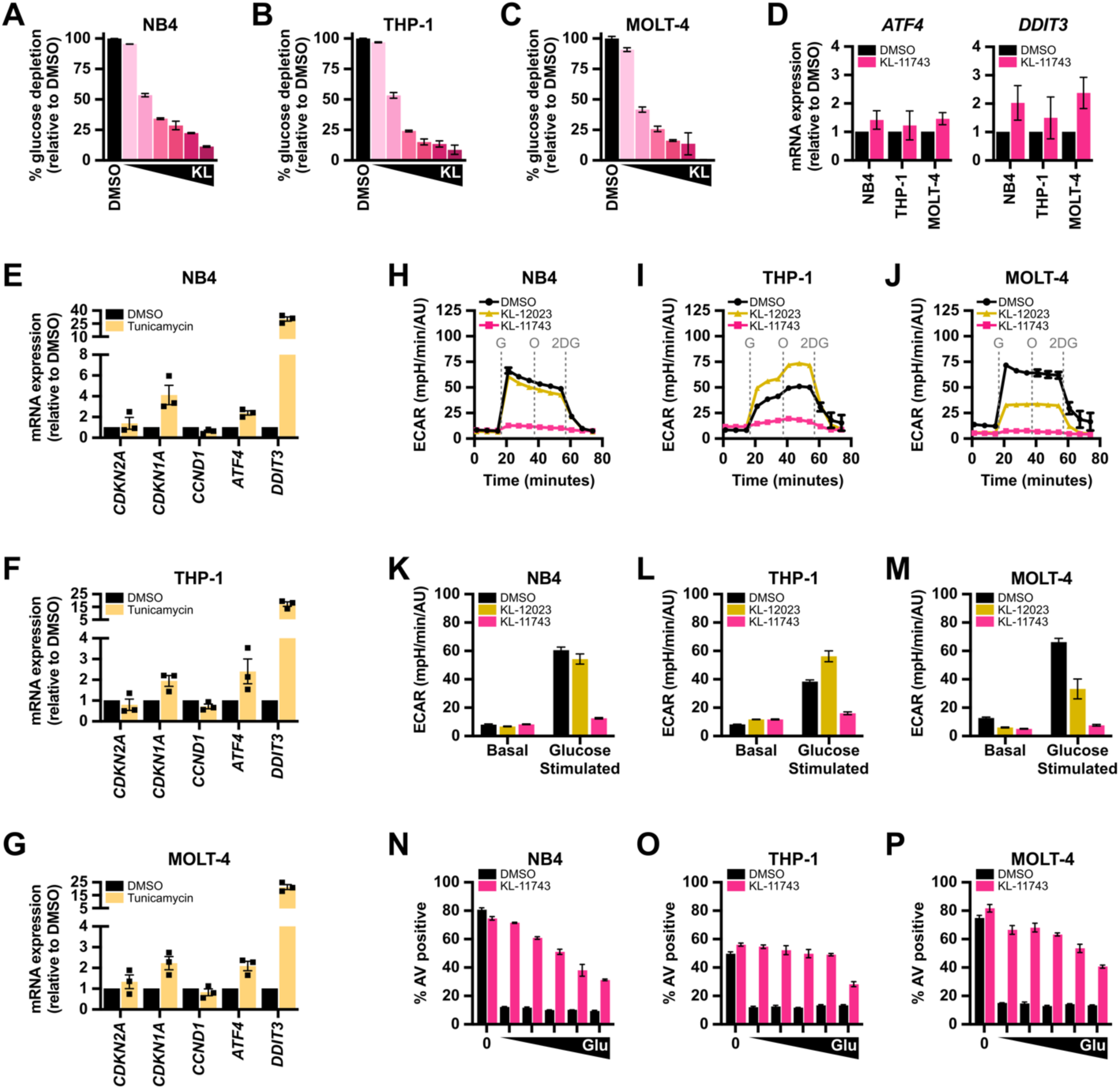
KL-11743, but not KL-12023, inhibits glucose uptake and glycolysis in hematological cells. **(A–C)** NB4, THP-1, and MOLT4 were treated with DMSO or KL-11743 (10, 100, 250, 500, 1000, 1500 nM) for 24 h. Different ranges were tested for each serum lot. Glucose concentrations from the cultured media were determined and presented as % glucose uptake relative to DMSO. Error bars are the SEM. **(D)** NB4, THP-1, and MOLT-4 were treated with DMSO or KL-11743 (500 nM) for 8 h, and total RNA was harvested. The fold change of transcripts for *ATF4* and *DDIT3* (CHOP) was determined by real-time qPCR. Expression was normalized against *18S*. **(E–G)** NB4, THP-1, and MOLT-4 were treated with DMSO or tunicamycin (100 ng/ml) for 8 h, and the indicated genes were determined by real-time qPCR. Expression was normalized against *18S*. Data are presented as the mean of 3 replicated experiments ± SEM. **(H–J)** NB4, THP-1, and MOLT4 were treated with DMSO, KL-11743 (500 nM), or KL-12023 (500 nM) for 24 h and ECAR was measured following an Agilent XF Glycolysis Stress Test. G: glucose (10 mM), O: oligomycin (1 μM), 2DG: 2-deoxy-D-glucose (50 mM). Error bars are the SEM. **(K–M)** Data from *H–J* presented as basal and glucose-stimulated ECAR. Error bars are the SEM. **(N–P)** NB4, THP-1, and MOLT4 were cultured in serum-free media for 24 h, then supplemented with glucose (0, 0.1, 0.5, 1, 5, 10 mM) ± KL-11743 (500 nM) for 24 h. Apoptosis was measured by AV labeling and flow cytometry. Error bars are the SEM.

**Supplemental Figure 2.**
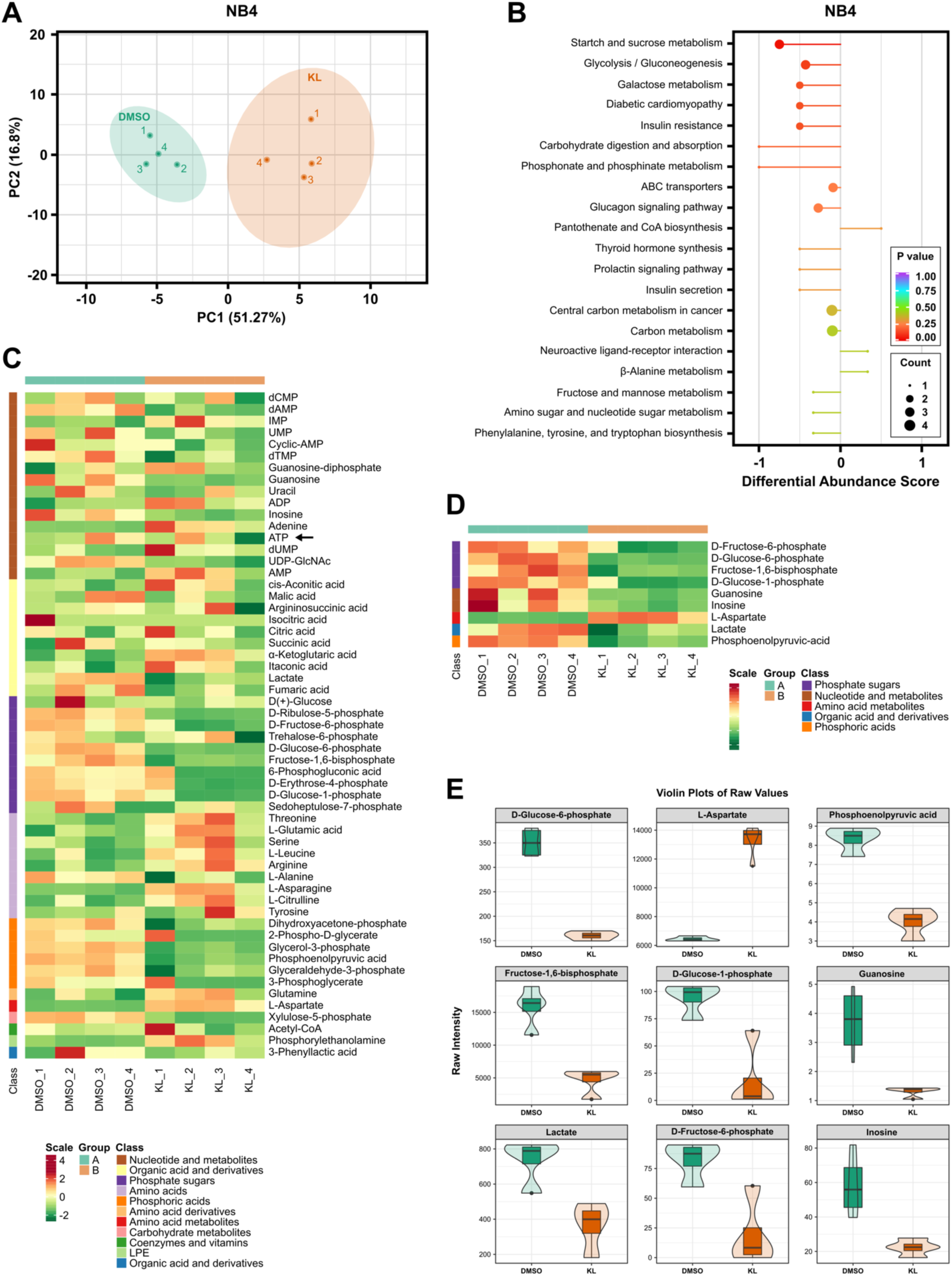
Targeted metabolomics of NB4 treated with KL-11743. **(A)** NB4 cells were treated with DMSO or KL-11743 (500 nM) for 24 h. Samples were subjected to LC-MS/MS and PCA detecting 57/68 energy related metabolites within the MetWareBio Energy Metabolism module. Unsupervised PCA was performed using prcomp within R; data were unit variance scaled before unsupervised PCA. **(B)** KEGG pathway enrichment analysis was conducted based on the annotation results. The size of the dots in the figure represents the number of significantly different metabolites enriched in the corresponding pathway. The X-axis represents the Rich Factor and the Y-axis represents the pathway. The color of points reflects the p-value. The darker the red, the more significant the enrichment. The size of the dot represents the number of enriched differential metabolites. **(C)** Hierarchical Cluster Analysis was used to cluster the samples. X-axis indicates the sample name and the Y-axis are the metabolites. Group indicates sample groups. Z-Score indicates the relative quantification of each metabolite with red representing higher content and green representing lower content. Cluster analysis was performed on both metabolites (vertical cluster tree) and samples (horizontal cluster tree). Heatmap was drawn by R software Pheatmap package. **(D)** Heatmap of different metabolites. The X-axis shows the name of the samples, and the Y-axis shows the differential metabolites. Different colors in the heatmap represent the values obtained after normalization and reflect the level of relative quantification. The darker the red, the higher the quantification. In contrast, the darker the green, the lower the quantification. The colored bar on top depicts sample groups. **(E)** Violin plots display data distribution and probability density. X-axis refers to sample, and the Y-axis refers to content. The box in the middle represents the interquartile range, and the middle box represents the 95% confidence interval. The black horizontal line is the median, and the outer shape represents the distribution density of the data. The figure shows the result of the top 9 differentially expressed metabolites with the largest Log_2_FC value.

**Supplemental Figure 3.**
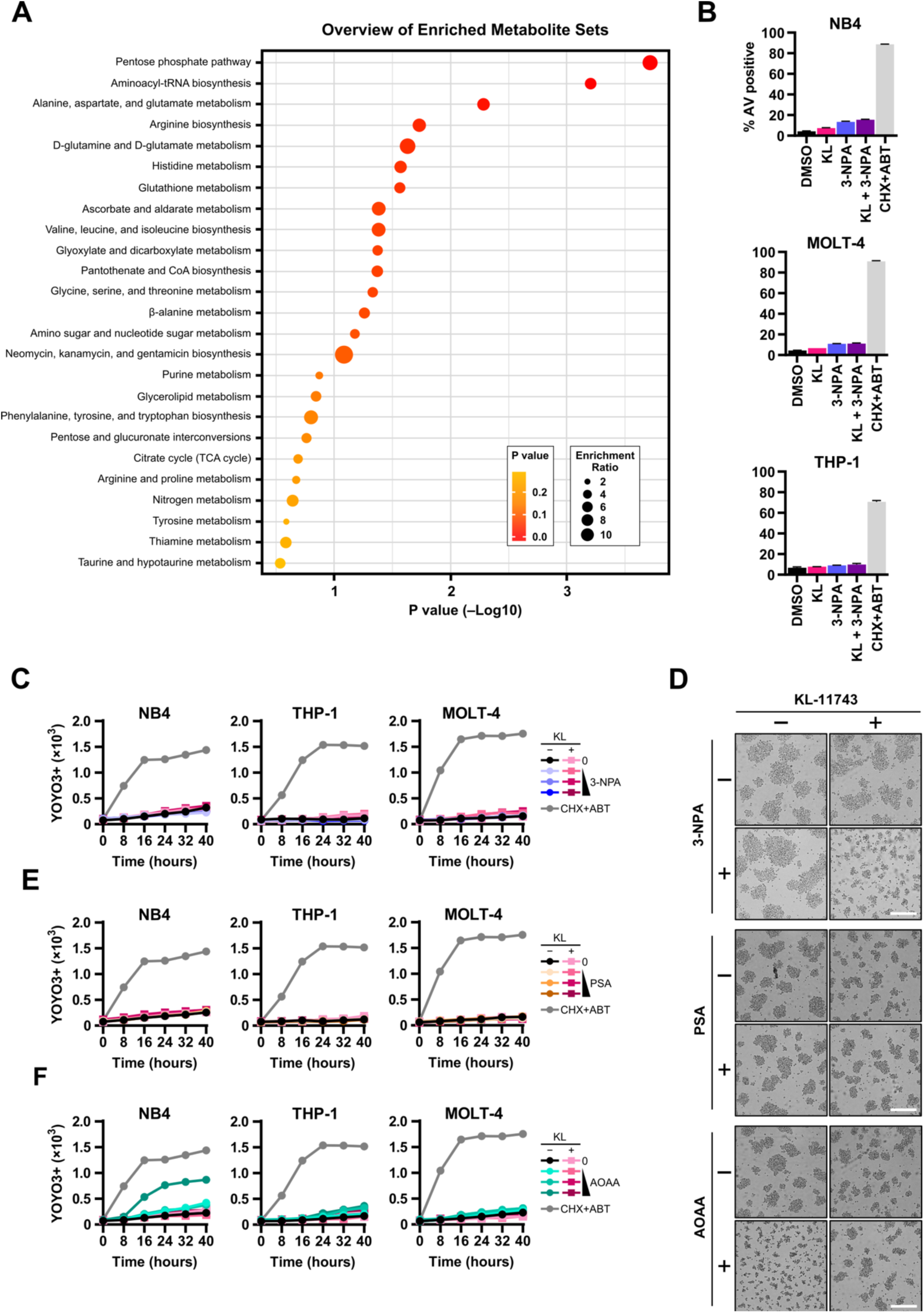
KL-11743 does not induce a dependency on the malate-aspartate shuttle. **(A)** Top 25 enriched metabolite sets in NB4 and MOLT-4 treated with KL-11743 (500 nM) for 24 h, compared to DMSO controls. P values were computed using an unpaired T test. **(B)** NB4, THP-1, and MOLT-4 were treated with DMSO, KL-11743 (KL, 500 nM) ± 3-NPA (1 mM) for 24 h. Apoptosis was measured by AV labeling and flow cytometry. CHX (50 μg/mL) + ABT-737 (1 μM) is a positive control for apoptosis. Data are presented as mean values of at least 3 replicates ± SEM. **(C)** NB4, THP-1, and MOLT-4 cells were treated with 3-NPA (0.5, 0.75, 1 mM) ± KL-11743 (500 nM), imaged every 8 h, and analyzed for YOYO3+ cells; the mean YOYO3+ events per image of 2 replicates is presented. CHX (50 μg/mL) + ABT-737 (1 μM) is a positive control for apoptosis. **(D)** Representative images of NB4 treated with KL-11743 (500 nM) and/or 3-NPA (1 mM), phenylsuccinic acid (PSA, 1 mM), or aminooxyacetic acid (AOAA, 1mM) for 24 h. Scale bar: 300 μm. **(E)** Same as *C*, but cells were treated with phenylsuccinic acid (PSA, 0.5, 0.75, 1 mM) ± KL-11743 (500 nM). **(F)** Same as *C*, but cells were treated with aminooxyacetic acid (AOAA, 0.5, 0.75, 1 mM) ± KL-11743 (500 nM).

**Supplemental Figure 4.**
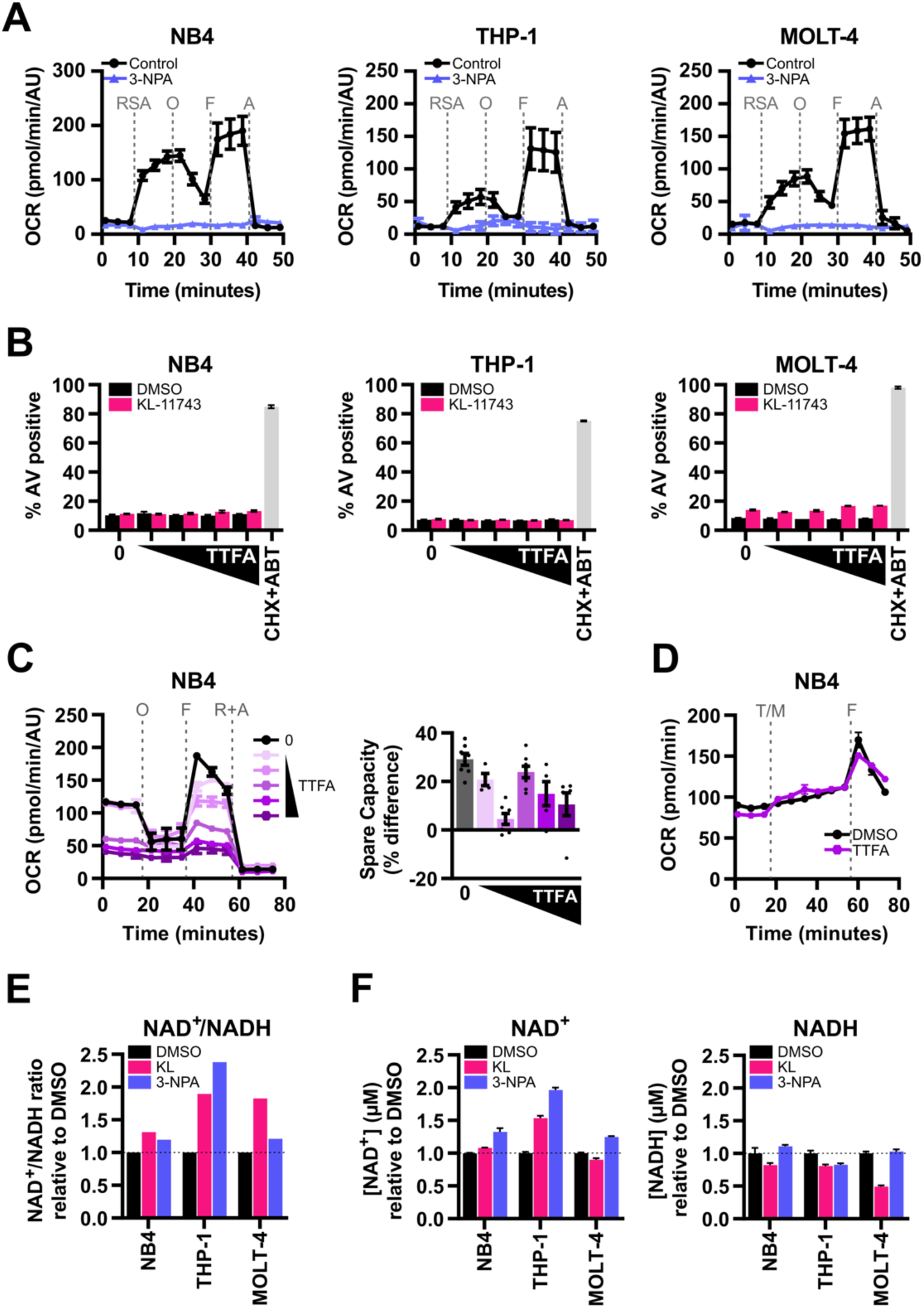
Disruption of TCA cycle and inhibition of glycolysis both alter NAD^+^/NADH redox. **(A)** CII analysis of NB4, THP-1, and MOLT-4 treated with 3-NPA (1 mM) for 24 h. OCR were measured by an Agilent XFe96 Analyzer during sequential administration of a combination of RSA: rotenone (1 μM) + succinic acid (10 mM) + ADP (4 mM), O: oligomycin (1 μM), F: FCCP (1 μM), and A: antimycin A (0.5 μM). **(B)** NB4, THP-1, and MOLT-4 were treated with DMSO, KL-11743 (500 nM), or TTFA (5, 10, 25, 50 μM) ± KL-11743 (500 nM) for 24 h. Apoptosis was measured by AV labeling and flow cytometry. CHX (50 μg/mL) + ABT-737 (1 μM) is a positive control for apoptosis. Data are the mean of 3 replicates ± SEM. **(C)** Left: NB4 were treated with TTFA (10, 20, 30, 40, 50 μM) for 24 h and OCR was measured following an Agilent XF Cell Mito Stress Test. O: oligomycin (1 μM), F: FCCP (1 μM), R+A: rotenone (0.5 μM) + antimycin A (0.5 μM). Right: Quantification of spare respiratory capacity calculated as the percent difference between basal and maximal respiration reads. **(D)** Changes in OCR of NB4 were measured using an Agilent XFe96 Analyzer after injection of TTFA (T, 50 μM) or XF RPMI medium (M) and FCCP (F, 1 μM). **(E-F)** NB4, THP-1, and MOLT-4 were treated with DMSO, KL-11743 (500 nM), or 3-NPA (250 μM) for 24 h before NAD^+^ and NADH concentrations were assessed. NAD^+^/NADH ratios in *E* were determined using average NAD^+^ and NADH concentrations from *F*. Data are displayed as mean values of 3–6 technical replicates ± SEM.

**Supplemental Figure 5.**
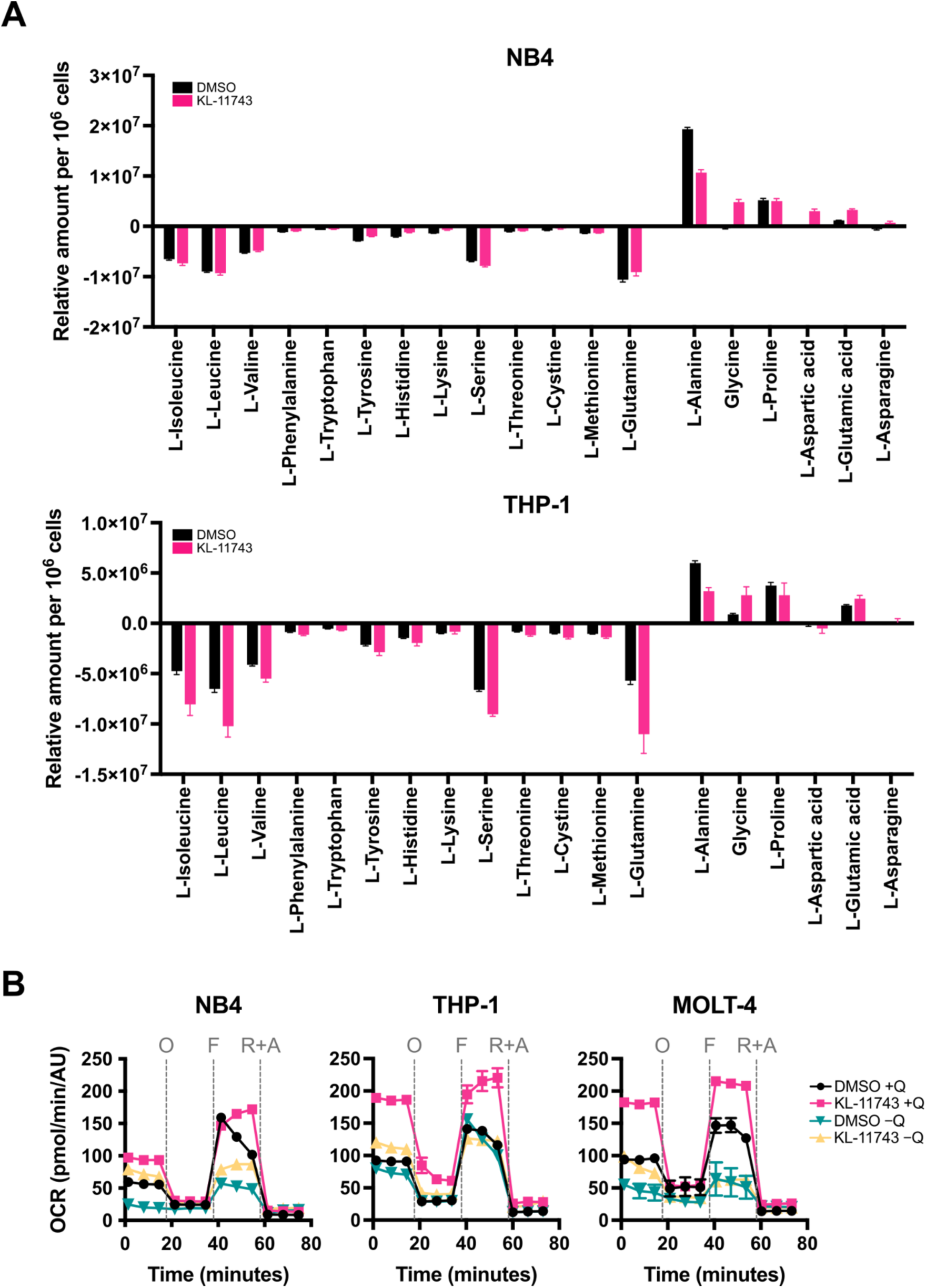
KL-11743 shifts amino acid metabolism. **(A)** NB4 and THP-1 were treated with DMSO or KL-11743 (500 nM) for 24 h, followed by YSI media analysis of all amino acids. Average rates are graphed ± SEM, with negative values indicating consumption and positive values indicating secretion. **(B)** NB4, THP-1, and MOLT-4 were treated with DMSO or KL-11743 (500nM) for 24 h prior to changing them into glutamine-free ((−)Q) media for 2 h. OCR was measured using an Agilent XF Cell Mito Stress Test. O: oligomycin (1 μM), F: FCCP (1 μM), R+A: rotenone (0.5 μM) + antimycin A (0.5 μM).

**Supplemental Figure 6.**
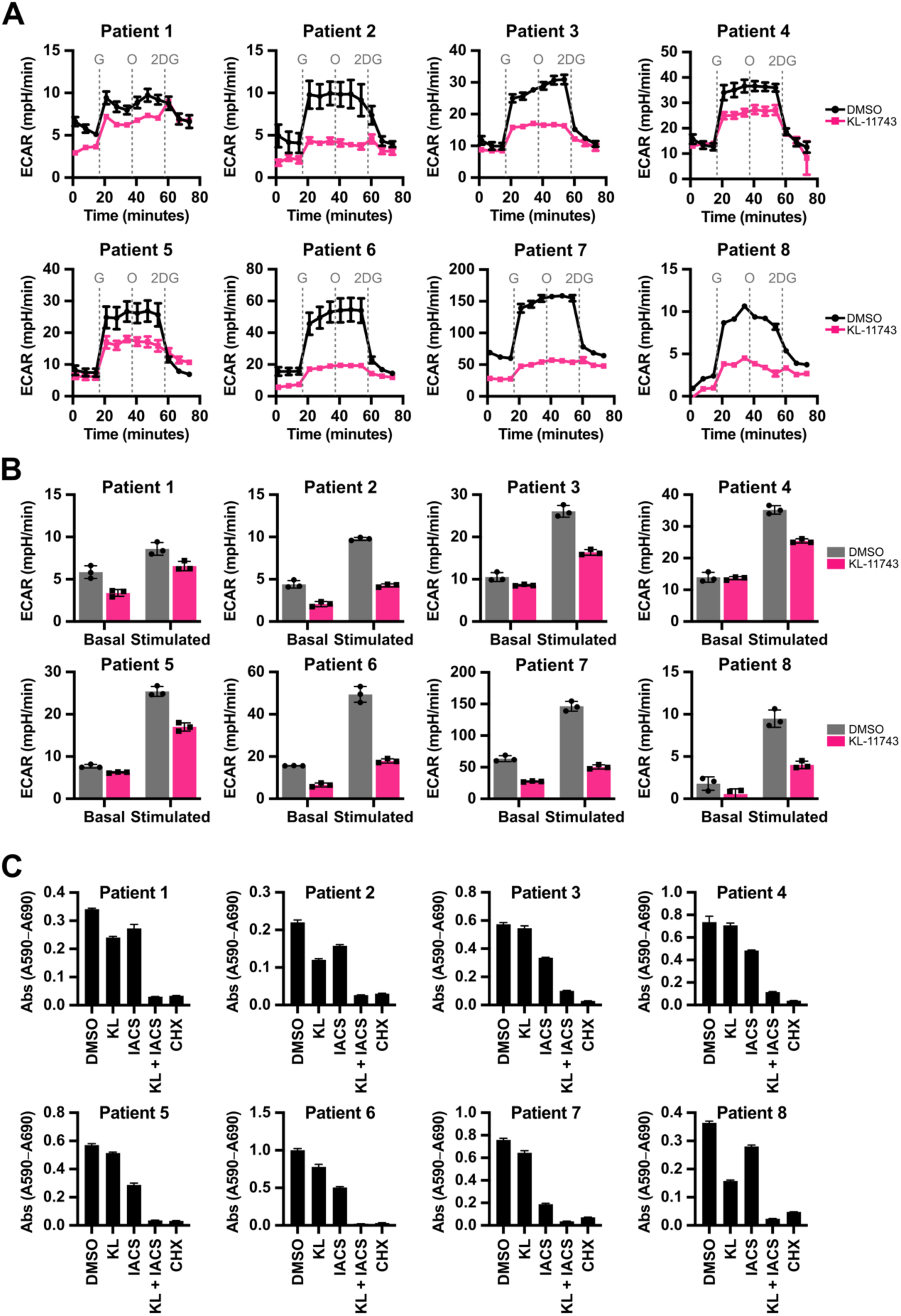
AML patient mutations and expanded clinical data. **(A)** Primary cells from AML patients 1–8 were treated with DMSO (0.1%) or KL-11743 (500 nM) for 24 h, and ECAR was measured following an Agilent XF Glycolysis Stress Test. G: glucose (10 mM), O: oligomycin (1 μM), 2DG: 2-deoxy-D-glucose (50 mM). All data are presented as mean values of at least 3 technical replicates ± SEM. **(B)** Data from *A* presented as basal and glucose-stimulated ECAR. Data are presented as the mean of 3 technical replicates, averaged over the time points before or after glucose injection, ± SEM. **(C)** Primary cells from AML patients 1–8 were treated with DMSO (0.1%), KL-11743 (500 nM), IACS (10 nM), CHX (50 μg/mL), or indicated combination for 24 h before viability was assessed by measuring the absorbance at 590 nm minus the reference absorbance at 690 nm. DMSO and CHX are the negative and positive cell death controls, respectively. All data are displayed as mean values of at least 6 technical replicates ± SEM.

